# Sedimentation and drag in drifting macrophytes and plastic objects: A model

**DOI:** 10.1101/2025.04.29.651208

**Authors:** Friederike Gronwald, Florian Weinberger, Tjeerd J. Bouma, Rolf Karez

## Abstract

Predicting macroalgal sedimentation and drag sensitivity is essential for ecological and geochemical modeling, and for optimizing seaweed cultivation. However, despite the diversity of macrophyte forms, models incorporating their specific morphology and hydrodynamic effects are largely lacking. To develop a broadly applicable model, we tested whether the drag response of diverse macrophyte morphologies and plastic objects can be accurately predicted by approximating them as ellipsoids and accounting for their specific shapes. A set of simple shape descriptors (wet weight, volume, thallus thickness, thallus projection area) and an empirical solution for the drag equation, enabled relative accurate predictions of the sinking velocity for 26 morphologically diverse macroalgae species, as well as the eelgrass Zostera marina, another major source of drifting biomass in many shallow seas. Additionally, we identified a second simpler empirical solution that incorporates shape and, while slightly less accurate, can be applied to a broader range of particles, including plastics.

Graphical abstract

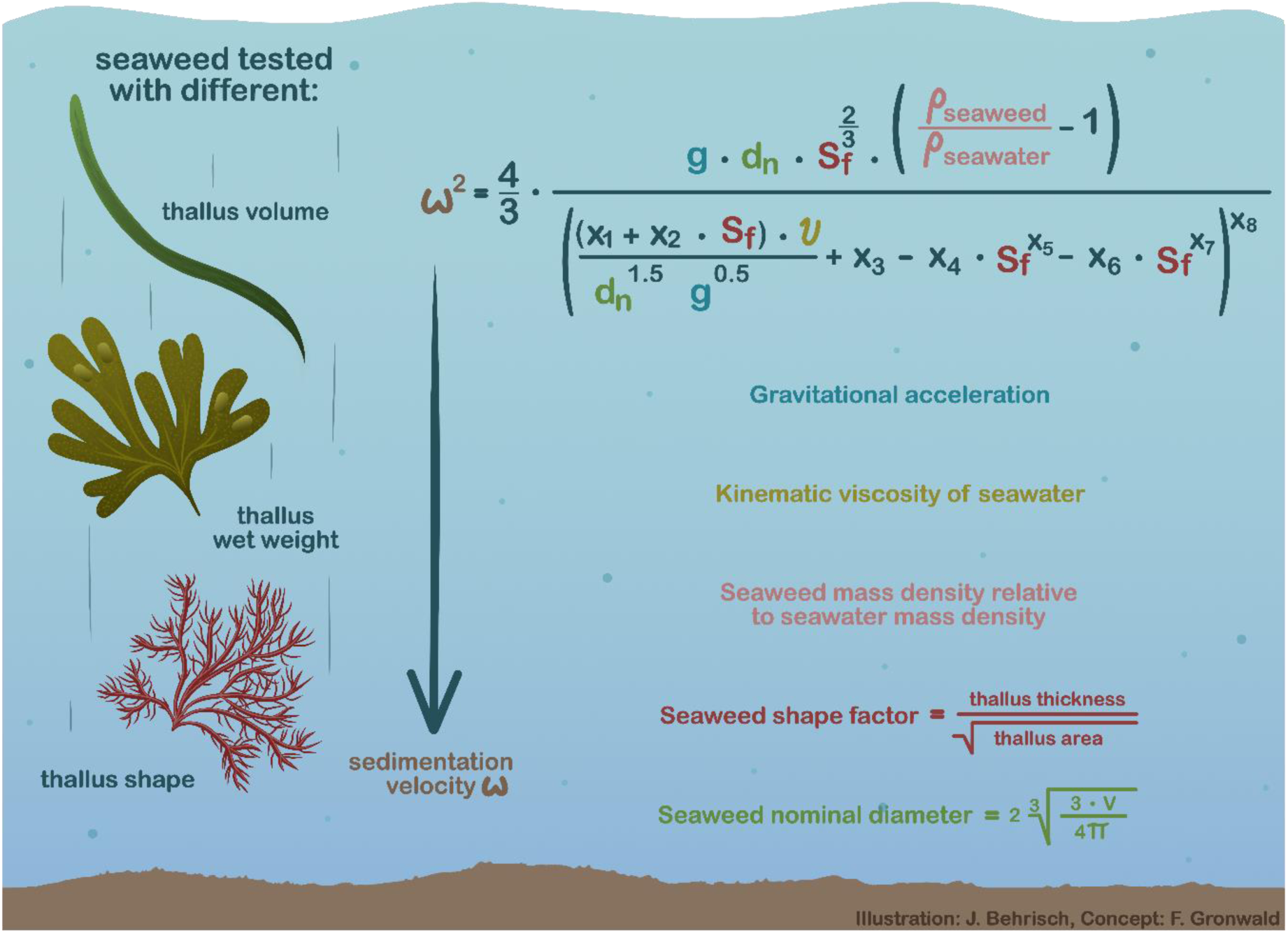

## Introduction

Drifting macroalgal blooms are a global problem, with some of the largest blooms being caused by floating species such as *Sargassum natans* and *S. fluitans* in the Great Atlantic *Sargassum* Belt or *Ulva prolifera* in the Yellow Sea (Smetacek and Zingone 2013). When blooms of that magnitude beach, they often cause severe problems for the coastal communities and environment (Zhang 2019, Bartlett and Elmer 2021). In the SW Baltic Sea the macroalgae blooms are of a smaller scale, but more diverse than in many other environments (Weinberger et al. 2021). These blooms can be dominated by a single species, but more often several species are found blooming together, and in most blooms in the SW Baltic, the eelgrass *Zostera marina* (littered leaves, fragments and whole specimens) also constitutes a significant part of the biomass (Weinberger et al. 2020). The ways in which hydrodynamic factors affect the species composition of macroalgal blooms are still poorly known. Unattached macrophytic biomass in the Baltic can be floating (Rothäusler et al. 2015), or negatively buoyant, but drifting (Bonsdorff 1992, Weinberger et al. 2008). Problematic blooms are often found in sheltered, relatively shallow waters, where they may degrade and cause nuisance and environmental problems locally (Mossbauer et al. 2012, Weinberger et al. 2020, 2021).

Accumulations of algal biomass can have great impact on the environment in deeper waters as well (Vahteri et al. 2000). It has been proposed that significant portions of biomass produced in photic coastal waters may even reach deep anoxic zones of the ocean through drift and sedimentation, where they could then provide an important component of global carbon sequestration (Krause-Jensen and Duarte 2016, Ortega et al. 2019, Kokubu et al. 2019). Such transport of particulate biomass over long distances obviously requires drifting velocities that exceed the speed of biomass degradation during the transport process. Possible maximum drift velocities of seaweeds are typically predicted using Lagrangian particle transport models (Rothäusler et al. 2015, Kwon et al. 2019, Garbossa et al. 2021, Zhou et al. 2021). These models do usually not consider the specific resistance of seaweeds to drag, as suitable generalized descriptors for predicting the hydrodynamic drifting behavior of seaweeds are so far largely missing.

The lack of models that can accurately describe the drifting and sinking behavior of seaweeds also affects the development and design of land-based seaweed aquaculture systems, which target the cultivation of unattached macroalgae. One important goal of such systems is to achieve maximally homogenous exposure of the cultivated organisms to sun light and nutrients with a minimum of energy investment (Sahoo and Yarish 2005). Typical solutions are raceways (Mata et al. 2003), aerated tank (Israel et al. 2005) or pond systems (Msuya and Neori 2008) or bioreactors (Savvashe et al. 2021) that are usually developed based on trial and error, as the drifting and sedimentation behavior of different seaweed species is difficult to predict.

In addtion to macroalgae and eelgrass, plastic particles represent another large group of drifting particles in the ocean (Eriksen et al. 2014). These particles can become problematic for example by remaining within the ecosystem and slowly degrading (Gewert et al. 2015), being ingested by marine organisms as microplastic (Galloway et al. 2017), or accumulating on beaches (Barnes et al. 2009). Plastic litter, to some degree, resembles seaweeds as it exhibits a similar diversity of shapes and sizes. These similarities suggest that a model developed to predict sedimentation velocities of seaweeds may also be applicable for plastic particles of similar size.

### Established approaches for predicting sedimentation velocity

As predicted by Stokes (1851), the sedimentation of particles in water is driven by the gravitational acceleration g. It also depends on the particle buoyancy, i.e. the particle mass density *ρ* relative to the mass density of the seawater *ρ*_sw_. Other factors that determine the sedimentation velocity ω are the diameter d of the particle and the drag coefficient C_D_. C_D_ is a dimensionless indicator of the resistance of a particle to current. C_D_ is not a constant, but varies as a function of flow speed, flow direction, object position, object size, fluid density and the kinematic viscosity of the medium. ω may be calculated for spherical particles as

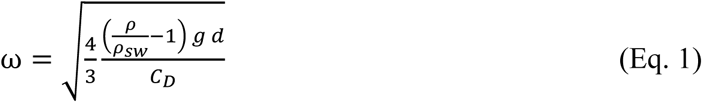

Equation 1 is based on the assumption that the sedimentation velocity can be calculated by equating the effective weight force with the drag force (Riazi and Türker 2019). For non-spherical particles, the nominal diameter d_n_ – describing the diameter of a sphere with the same volume as the particle – can be used instead of the diameter d. However, exact analytical solutions of Eq. 1 only exist for spheres (Stokes 1851) and spheroids (Oseen 1927) in laminar flows, because turbulence on the particle surface strongly influences C_D_ and irregularly shaped particles must be expected to generate turbulences that are virtually impossible to predict. To resolve this problem, different empirical solutions of Eq. 1 that used shape factors to correct C_D_ for non-sphericity have been proposed for sediment grains (e.g., Swamee and Ojha 1991, Cheng 1997, She et al. 2005, Camenen 2007). Particle shape could be expressed in various ways, for example by use of sphericity factors that put the volume of a particle in relation with its surface area (Wadell 1935). Yet, exact measurements of the surface area of irregularly shaped particles are often hardly possible, which makes the application of sphericity factors to them difficult. The shape factor most commonly used instead is the Corey factor S_f_ (Komar and Reimers 1978), which expresses the deviation of particle shape from sphericity independent of its size as

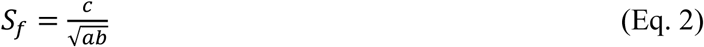

where a, b, and c are the diameters in the longest, the intermediate and the shortest mutually perpendicular axes of the particle, respectively. An improved empirical solution of Eq. 1 was recently proposed by Riazi and Türker (2019), who investigated the sedimentation behavior of sediment particles with diameters in the approximate range between 0.5 and 7 mm. The authors proposed to treat such particles as ellipsoids and introduced S_f_ raised to the power of 2/3 into Eq. 1 to account for their non-spherical shapes:

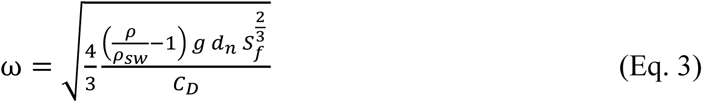

Riazi and Türker (2019) (Riazi and Türker 2019) further proposed to calculate the drag factor in Eq. 3 as

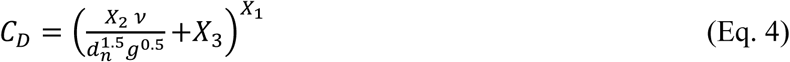

where ν is the kinematic viscosity of the medium, X_1_ is a dimensionless constant and X_2_ and X_3_ are also dimensionless constants that both depend on S_f_ and describe the behavior of the drag coefficient with respect to particle shape in laminar and turbulent flow conditions, respectively.

### The need to transfer these equations towards seaweed-like particles

Drifting macrophytes exhibit highly variable morphologies that range from unbranched filamentous forms over branched or unbranched leaf shapes to three-dimensionally branched or unbranched irregularly tangled forms (Fig. S4 to S10). Macrophytes are usually much larger and mechanically more flexible than sediment grains, which may also be expected to affect their response to drag. The question therefore arises whether generalized approaches to the determination of C_D_ and ω are possible that allow for the prediction of sedimentation velocities and drag across species without tedious experimental setups. We hypothesized that approximate predictions may be possible if drifting macroalgae and eelgrass litter are treated as ellipsoids, similar as proposed by Riazi and Türker (2019) for sediment particles.

The approximately ellipsoidal character of unbranched macrophytes is fairly obvious. However, in macrophytes exhibiting branching of several orders typically the longest, shortest, and middle perpendicular thallus axes cannot be readily identified and measured. It was therefore necessary to develop a suitable method for the determination of these axes and S_f_ in macrophytes. Sedimentation velocities of two different sample sets of seaweeds and eelgrass and a set of plastic objects, with variable particle properties were then measured. Measures obtained with seaweed sample set 1 allowed for the identification of an empirical solution for Eq. 4 and the resulting model was successfully tested with seaweed sample set 2. Subsequently, it was tested whether the optimal solution for seaweeds would be applicable to plastic particles.

## Material and methods

### Collection and maintenance of algae

This study includes 73 specimens of drifting marine macrophytes belonging to 26 different species (*Fucus vesiculosus, Fucus serratus, Chorda filum, Saccharina latissima, Gracilaria vermiculophylla, Ceramium virgatum, Vertebrata fucoides, Polysiphonia stricta, Spermothamnion repens, Ahnfeltia plicata, Furcellaria lumbricalis, Coccotylus truncatus, Delesseria sanguinea, Cladophora flexuosa, Cladophora* sp.*, Rhodomela confervoides, Pyropia leucosticta, Ulva clathrata, Ulva linza, Ulva compressa, Kornmannia leptoderma, Bryopsis hypnoides, Acrosiphonia centralis, Zostera marina*, *Ascophyllum nodosum* and *Ulva gigantea*). The specimens were selected to represent a wide range of morphologies (Fig. S4 to S10) and were taxonomically identified based on these traits, following Nielsen et al. (2023).

Ten of the specimens were collected from drift material in January 2023 at a location between Strande and Bülk light house (Kiel Fjord/Germany; 54°26’57.4”N 10°11’37.6”E). On the 26th of June 2024, 56 more specimens were collected at the same site and at two other locations in the Kiel Fjord area (Schilksee, 54°25’16.3”N 10°10’43.1”E and Mönkeberg, 54°21’20.92”N 10°10’41.97”E). Six specimens of *A. nodosum* and one of *U. gigantea* were collected in July 2024 from a beach at Yerseke/Netherlands (51°30’09.0”N, 4°02’39.7”E). Salinities and water temperatures at the collection sites and during subsequent maintenance and experiments were measured using a WTW Multi3630IDS conductometer. Prior to use, material collected in 2023 was maintained for 3 d at 13.6 °C in Baltic Sea seawater with a salinity of 18.8 psu, which was the salinity at the collection site. All subsequent measurements were conducted in sea water of the same salinity and temperature at the GEOMAR laboratories in Kiel. Material collected from the Kiel Fjord in 2024 was kept for 4 days in seawater with salinity 15.0 psu (mean salinity at the collection sites: 14.0 psu) and 16 °C (mean temperature at collection sites: 20 °C) and then packed in cooler boxes and transported within 7 h to the NIOZ laboratories at Yerseke in the Netherlands. There the material was maintained for 1 to 3 weeks at 18 °C in Oosterschelde water (salinity: 33 psu) diluted with tap water to a salinity of 14.4 psu. Material collected from the Oosterschelde was acclimatized at the NIOZ laboratories to a salinity of 15 psu by stepwise decrease of salinity by 4-5 psu every two to three days. Aeration was provided to all specimens during the maintenance and they were kept in artificial light (80 µmol photons m^−2^ s^−1^ for 12 h d^−1^). Water was exchanged every other day.

In addition to the seaweeds, 16 negatively buoyant plastic objects were tested, including eight circular foil cutouts (disks), three table tennis balls, two plastic nets and three rubber bands (Fig. S11). The foil disks were cut to different diameters and some were punched with different numbers of small holes. The name of the foil circles indicates both their diameter and perforation level. For example, “Disk 40-1” had a diameter of 40 mm and was unpunched, where “1” denotes unpunched, “2” partially punched, and “3” heavily punched, “4” extremely heavily punched. The three table tennis balls shared identical dimension and therefore had the same shape parameters. To increase their mass density air inside the balls was replaced with sea water, using an injection syringe. To increase the mass density further different nubers of glass beads (2 mm diameter) were pressed into the balls through small holes, which were then sealed with Scotch tape.

### Descriptors of thallus shape and density

Particle traits were determined for all specimens. To determine volume V a graduated cylinder measure of suitable size was filled with a defined volume of sea water and the specimen was completely immersed in this volume. Any air bubbles were carefully removed and the increase of volume was recorded as particle volume. Wet weights were measured after the macrophytes had been carefully blotted with paper. All measurements of volume and weight were repeated two times and in case of data divergency by more than 5 % three times. The particle mass density ρ could then be calculated as the ratio of mean blotting weight and mean volume. The nominal particle diameter d_n_ was also derived from the mean volume and determined as twice the radius of a sphere with the volume of the measured macrophyte:

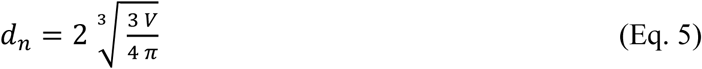

Projection images of all specimens were used to determine measures for the calculation of Corey shape factors. To generate such images the specimens were scanned on a flatbed scanner together with a size standard. The resulting images were analyzed using the Fiji software package (Schindelin 2012). In the case of specimens with flattened morphologies (*A. nodosum*, *F. vesiculosus*, *F. serratus*, *S. latissima* phylloid, *P. leucosticta, C. truncatus, U. compressa*, *U. gigantea, K. leptoderma, Z. marina* leaves) and also of the unbranched cylindrical *C. filum* the determined parameter was the thallus projection area (using the “adjust color threshold” and “measure particles” procedures implemented in Fiji). This projection area obviously represents the product of the longest and the intermediate perpendicular thallus axes a * b of flattened thalli (see Eq. 2). The shortest intermediate perpendicular axis c is in these cases equal to the thallus thickness. It could be subsequently calculated by division of the thallus volume V by the projection area a*b, because V = a*b*c. All remaining specimens exhibited three-dimensional morphologies, often with branches that partly overlapped and masked each other in projection images. In these cases, ten independent measurements of the width of youngest thallus branches were conducted and the mean was considered as a representative measure of c. Division of volume V by c allowed then to calculate corresponding values for a*b.

### Measurements of sinking velocity

Maximum sinking velocities in sea water were determined for all non-buoyant specimens directly before or after the determination of particle traits. After their release at the water surface, sinking particles initially accelerate until they reach their maximum sinking speed. It was therefore necessary to first determine a water depth below which a maximum sinking speed of the macrophytes could be assumed. It was found that a water depth of 15 cm is required for the acceleration phase. To verify this, the sinking speeds of 22 samples were measured in two different ways (Fig. S1). In both cases, the samples were released at the water surface and only after they had sunk to a depth of 15 cm was their speed measured during the further descent. In the first case, however, the sinking speed was only measured over the depth segment from 15 cm to 41 cm and in the second case over the depth segment from 15 cm to 76 cm. If the maximum speed is not reached at a depth of 15 cm and thus a further acceleration takes place below 15 cm, this should lead to higher mean sinking speeds for measurements at a greater depth (i.e. over a distance of 61 cm) than for measurements at a lesser depth (i.e. over a distance of 26 cm). However, a significant difference between the two measurements was only observed for two specimens (Welch corrected t-tests, p = 0.05), and in only one of these cases was the measured velocity higher when the measuring distance was longer. Moreover, a non-significantly higher mean sinking velocity was found for 13 specimens when measured over the shorter depth distance, and only for 7 specimens over the longer distance.

Therefore, sedimentation velocities were measured in such a way that the specimens were submerged just below the water surface and released, so that they could accelerate from the water surface to a depth of 15 cm before the velocity was measured with a stop-watch until the specimens made first contact with the bottom. In 2024 the measurements with small algae were carried out in a glass cylinder (diameter 40 cm), which allowed for lateral observation. In this case, the length of the fall distance was 87 cm and the monitoring distance was 72 cm. For larger algae, a plastic barrel was used, in which an underwater video camera and corresponding depth marks allowed for observation of the algae reaching the water depth of 15 cm. In these cases, the monitoring distance was 61 cm. All measurements were repeated five times. In 2023 sedimentation velocities were measured in an aquarium that was filled to a height of 26 cm with sea water and the monitoring distance was 11 cm. The macrophytes were filmed outside the aquarium, with a ruler attached to the window pane, and sedimentation times were determined by single frame analysis. These measurements were repeated 7 times. The plastic particles were measured in a plastic cylinder (diameter: 19 cm) with a fall distance of 48 cm and a monitoring distance of 33 cm.

Water temperature and salinity in the measuring vessels changed slightly from day to day. Both parameters were therefore determined repeatedly between the speed measurements, in order to derive mass density ρ_H2O_ and kinematic viscosity ν (Sharqawy et al. 2012). These parameters are listed in Tab. 1

**Table 1:**
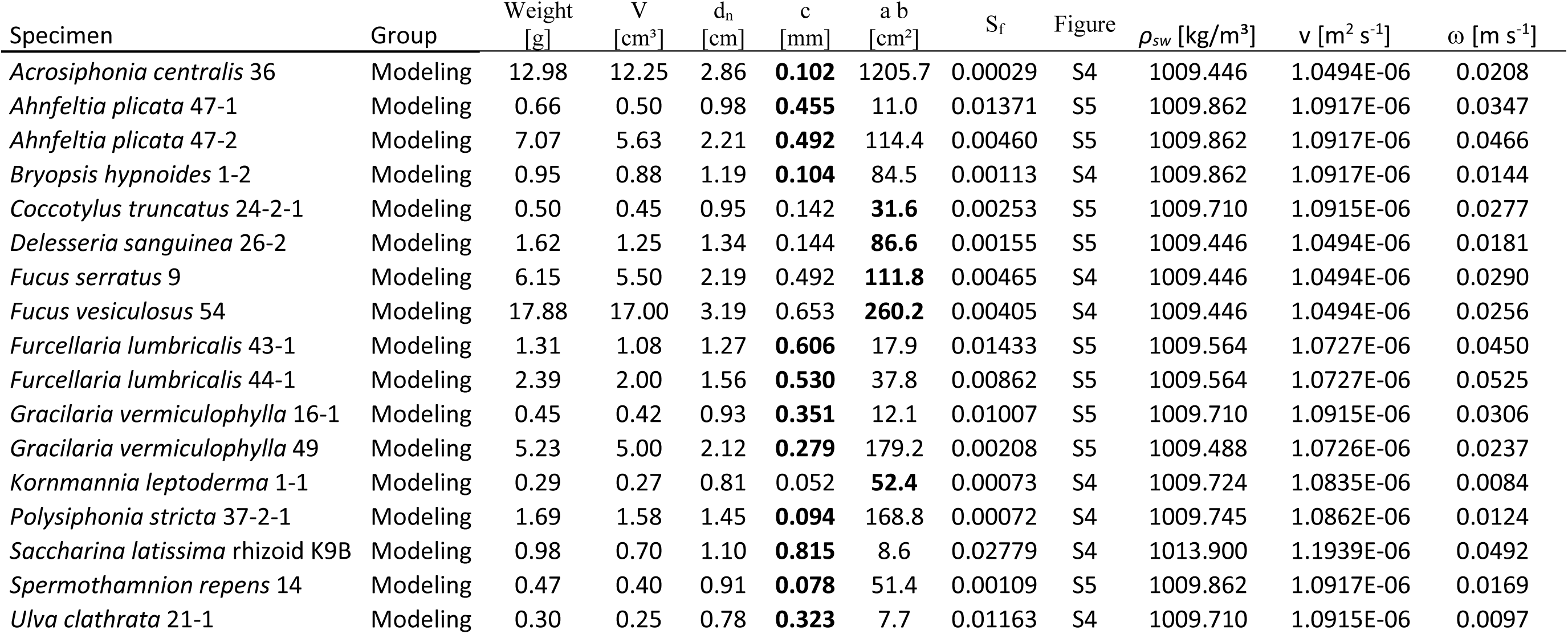
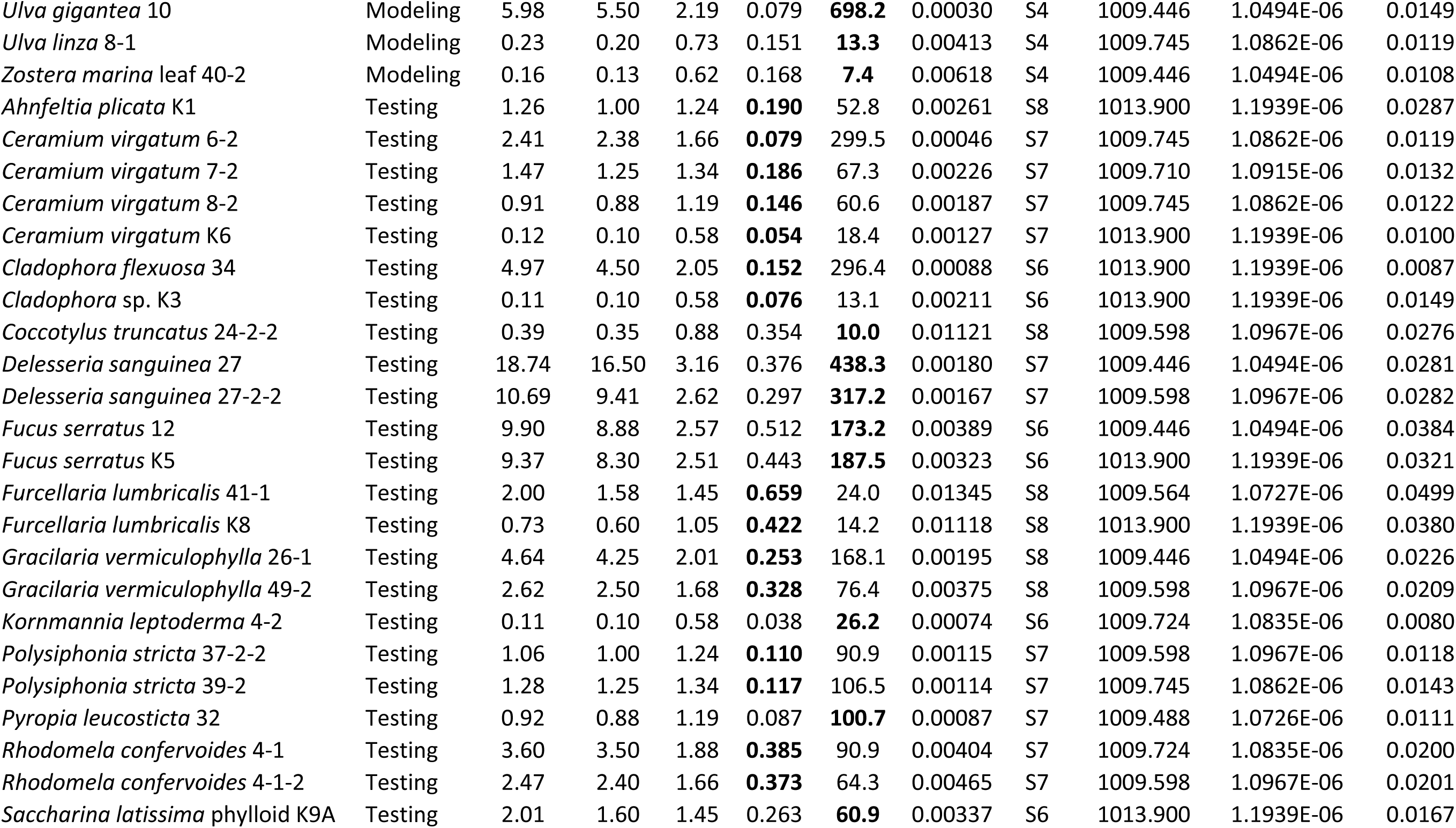
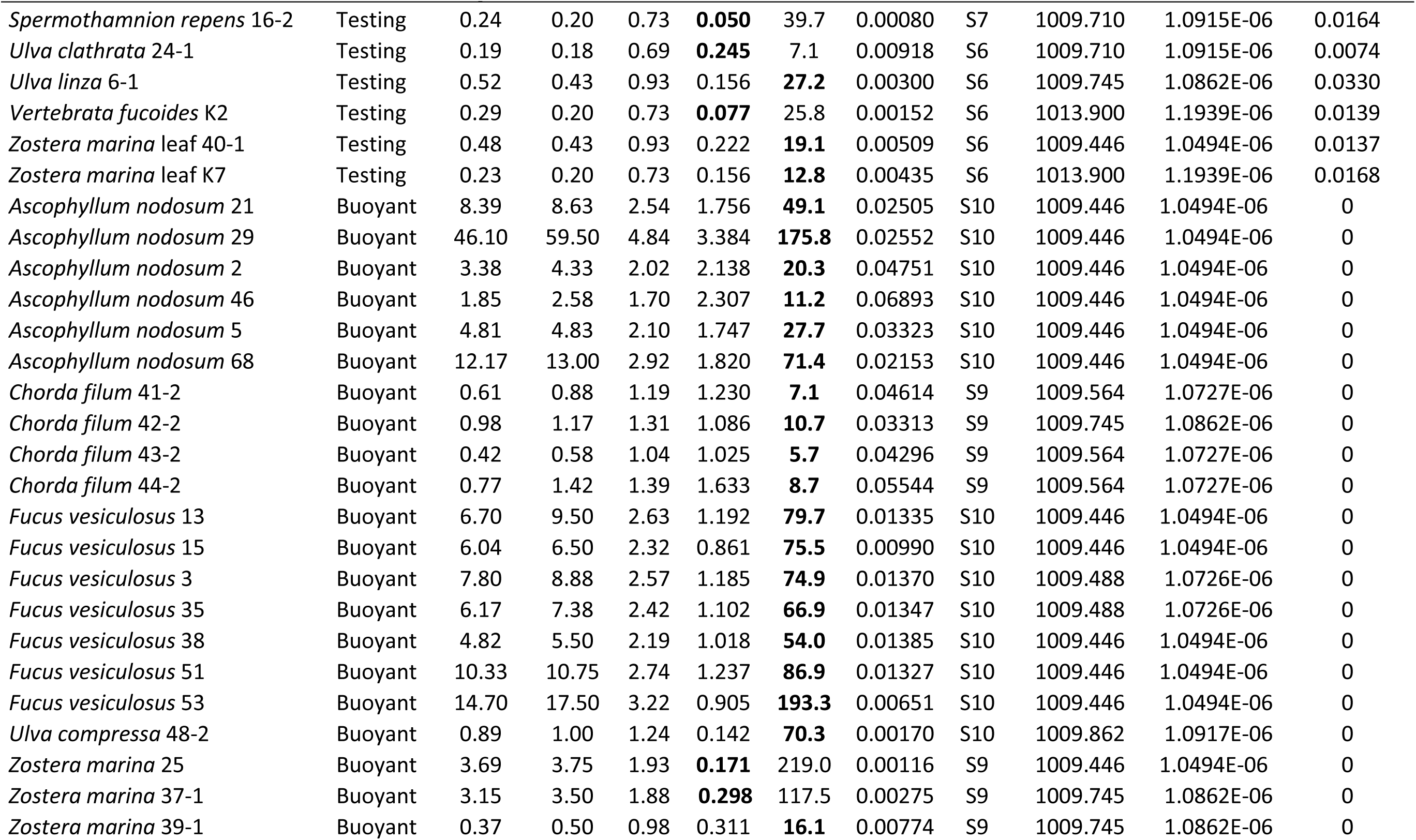
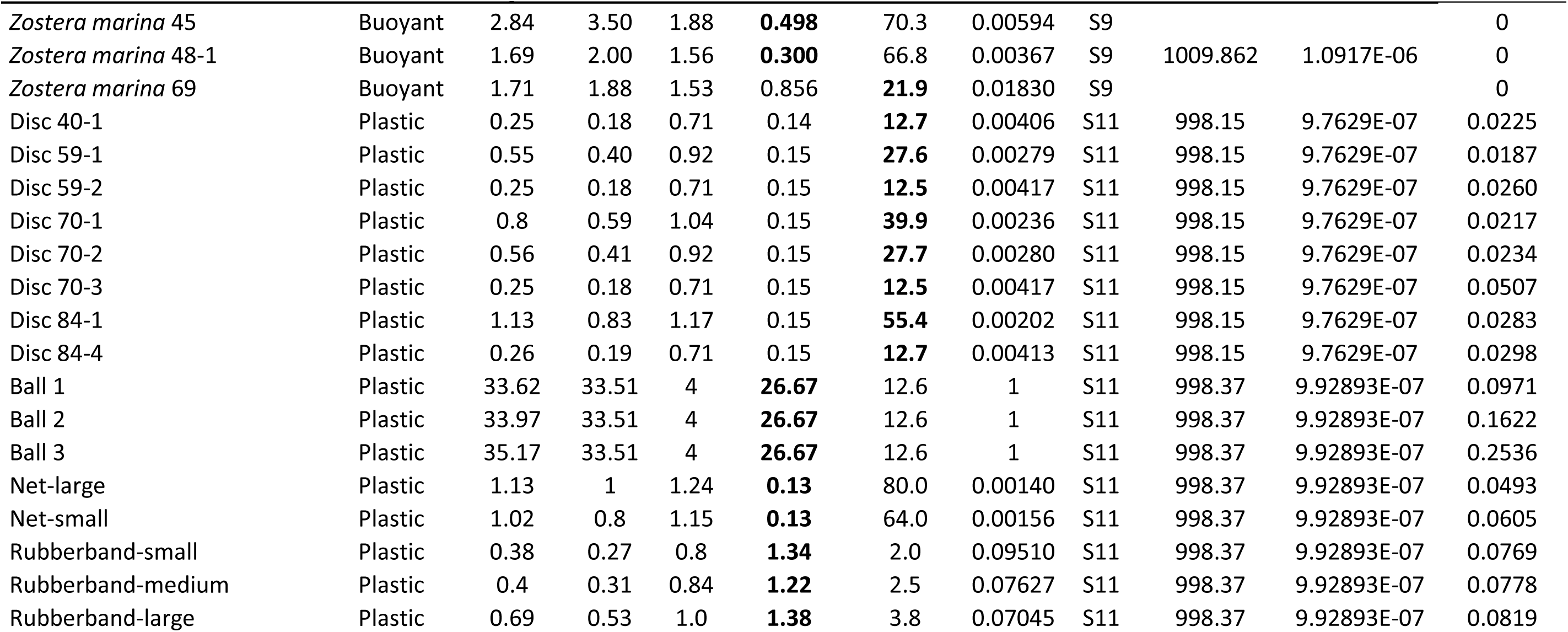
Shape descriptors of the investigated specimens. Non-buoyant specimens were grouped and used either for development or for testing of the sedimentation models, while buoyant specimens could obviously not be used for this purpose. Weight is the blotting weight [g], V is the volume [cm³], d_n_ [cm] is the nominal diameter. c [mm], a and b [cm] are thallus diameters in the shortest, longest and intermediate mutually perpendicular axes, respectively, that were used to calculate the Corey shape factor S_f_ (see Eq. 2). Bold values listed under c and a b were measured directly, while non-bold values listed under c and a b were determined by division of V by the bold value. Figure refers to pictures in the online supplement. ρ_H2O_ and ν are seawater density and kinematic viscosity during measurements of the mean sedimentation velocity ω.

### Statistics and model development

Data were visualized and correlations were analyzed using the Prism9 software package (GraphPad, Boston, USA). Differences in sedimentation velocities among specimens or groups of specimens were detected by Kruskal-Wallis-ANOVA and Sidak-corrected pairwise Dunn-tests using the same software. All other statistical treatment was conducted in R version 4.2.2. The sedimentation velocity equation (see Eq. 3 and 4) was optimized for the particles using genetic algorithm as implemented in R library GA (Scrucca 2013). For this purpose, the data sets of sedimenting macrophytes were divided into two groups. The first group was selected to include a wide range of different morphologies and particle properties, and this group was used for the actual modeling. The employed genetic algorithm started with an initial population of 5000 randomly generated individuals, with a crossover probability of 0.8, a mutation probability of 0.1 and an elitism of 250. The optimization process continued for maximally 500 000 generations and ended when no convergence toward a lower mean square error was obtained over 5000 generations. The accuracy of the resulting equations was then verified, using the second group of seaweed data sets as well as the plastic particle data set.

## Results

### Particle traits

Particle traits of all investigated specimens are summarized in Tab.1, for images of all specimens see Figs. S4 to S11. Nominal diameters of these specimens ranged from 0.576 cm to 4.844 cm, volumes from 0.1 cm³ to 59.5 cm³ and blotting weights from 0.11 g to 46.10 g. The shape factor S_f_ varied beween 0.00029 and 0.0689 (Fig. 1), with the exception of some of the plastic particles with higher shape factors, especially the table tennis balls, which had an S_f_ of 1 due to their perfectly round shape. As to be expected, S_f_ was particularly low in specimens with pronounced thin and flattened morphologies, such as *U. gigantea*, *K. leptoderma,* or *P. leucosticta*, but also in specimens characterized by very thin and dense lateral branches, such as *A. centralis*, *C. flexuosa* or *S. repens*. S_f_ was the highest in macrophyte specimens exhibiting relatively thick branched or unbranched morphologies, such as *C. filum*, *A. nodosum* or *S. latissima* rhizoid. A particularly large variability of S_f_ was detected for *Z. marina*, corresponding with the circumstance that the investigated specimens were morphologically variable and included littered leafs, as well as whole plants bearing or not bearing rhizoid or inflorescence. In *F. vesiculosus* floating specimens – characterized by presence of gas-filled vesicles – exhibited a larger S_f_ than non-floating specimens. A trend towards larger S_f_ in floating specimens gets also apparent if different species are compared: in none of the negatively buoyant seaweed specimens was S_f_ larger than 0.0278 and in none of the buoyant specimens smaller than 0.00116 (Fig.1).

**Fig. 1:**
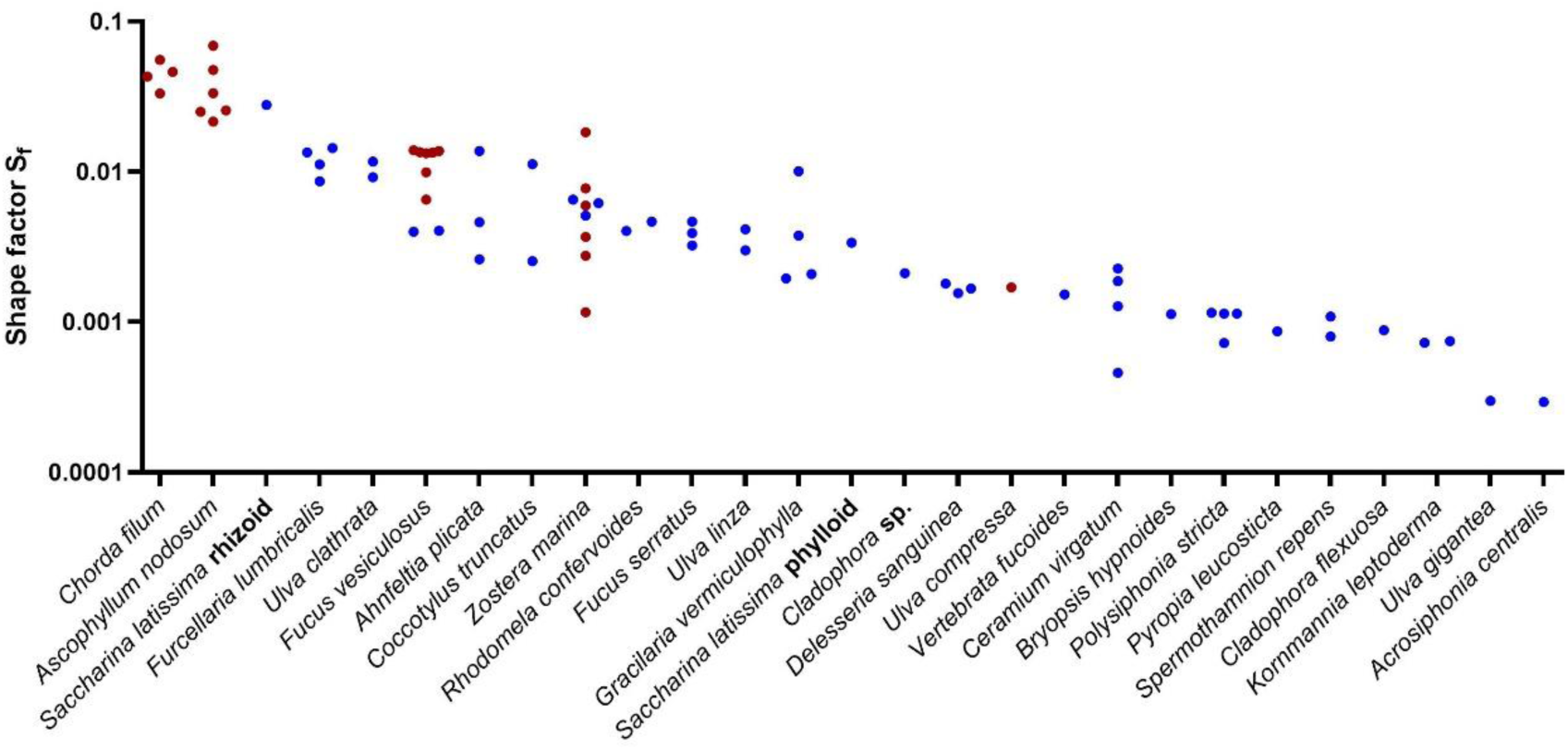
Shape factors of 73 specimens of drifting marine macrophytes. Red dots indicate positively buoyant (floating) specimens, blue dots negatively buoyant (sinking) specimens.

### Sedimentation velocities

Alltogether 49 specimens of macrophytes were negatively buoyant and could be used to determine sedimentation velocities. This group only included single leafs and no whole plants of *Z. marina*. The sedimentation speed of the 49 specimens varied over a relatively wide range (Fig. 2). The *S. latissima* phylloid sank considerably slower than the *S. latissima* rhizoid and one of the two specimens of *U. linza* – containing some air entrapped in its tubular thallus – sank less than half as fast as the second, which contained no air. In most other cases specimens belonging to the same species sank with more similar velocities. Particularly fast velocities were recorded for the rhizoid of *S. latissima*, as well as for the specimens of *F. lumbricalis* and one of the specimens of *A. plicata*. Particularly low velocities were observed with specimens of *K. leptoderma*, *C. flexuosa* and *U. clathrata*, which are all members of the Ulvophyceae. Indeed, green algal specimens generally exhibited significantly lower mean sinking velocities (Kruskal-Wallis-ANOVA; χ² = 13.91; df = 3; p = 0.003) than brown algal specimens (Dunn-test; p = 0.0069) or red algal specimens (p = 0.0441), while other differences among major taxonomic groups were insignificant (p > 0.05). Further, specimens exhibiting flattened branched morphologies (i.e., fucoids, *C. truncatus*, *D. sanguinea*) sank significantly faster (Kruskal-Wallis-ANOVA; χ² = 8.476; df = 2; p = 0.014) than specimens exhibiting flattened unbranched morphologies (i.e., *P. leucosticta*, *U. gigantea*, *U. linza*, *K. leptoderma*, *S. latissima* phylloid, *Zostera* leaves; Dunn-test, p = 0.0112). No significant difference was detected between sinking velocities of non-flattened and either flattened unbranched or flattened branched morphologies (p > 0.05).

**Fig. 2:**
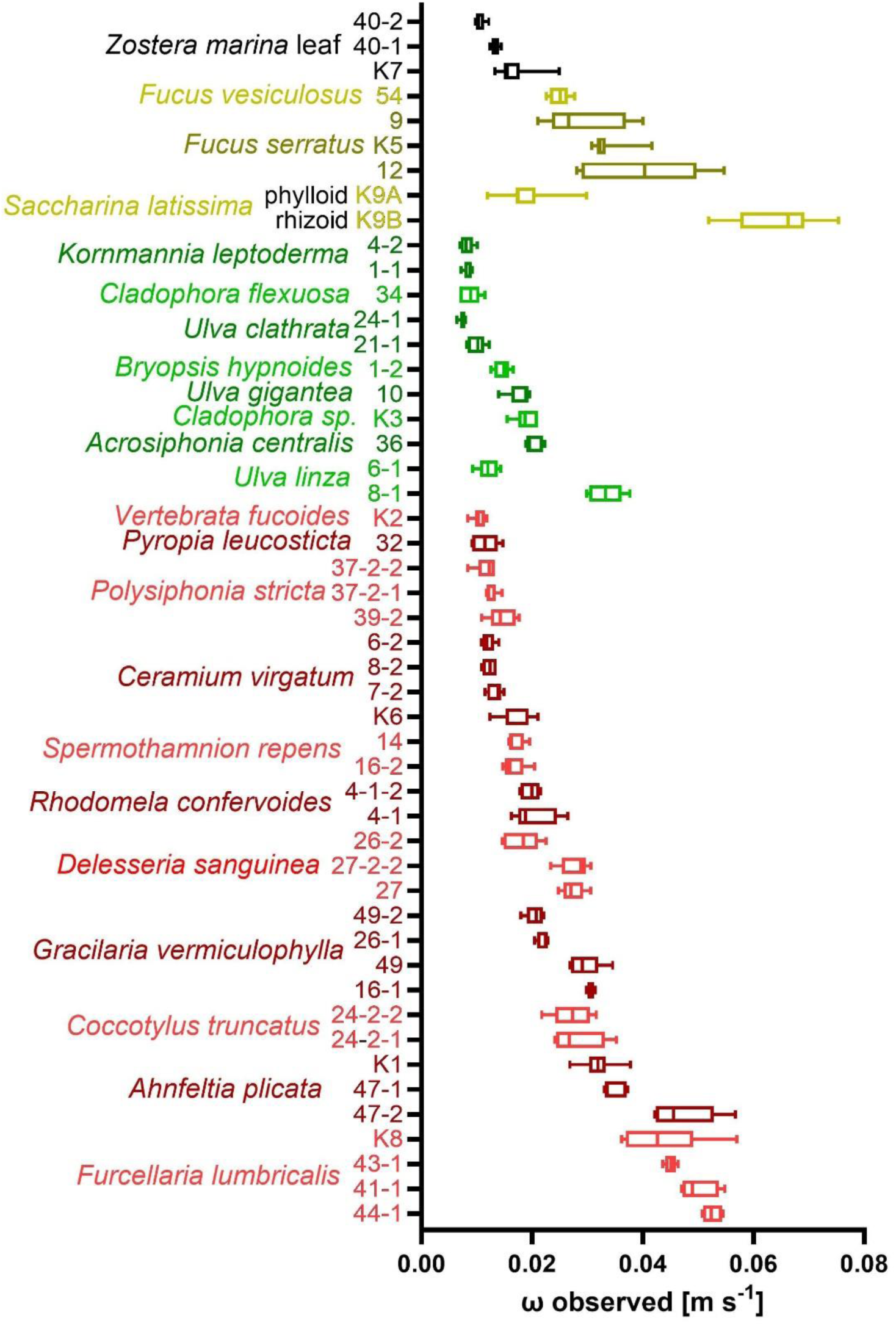
Sedimentation velocity ω of 49 specimens of macroalgae. Brown bars: Phaeophyceae, red bars: Rhodophyceae, green bars: Ulvophyceae and black bars: eelgrass. Median ± quartiles (n = 5; n = 7 for specimens with numbers beginning with „K“).

The sedimentation velocities were found to correlate significantly and non-linearly with both the nominal particle diameter of the macrophytes and their mass density relative to the density of the water at the time of measurement (Fig. 3). However, the strongest correlation was observed between the velocity and the shape factor of the particles.

**Figure 3:**
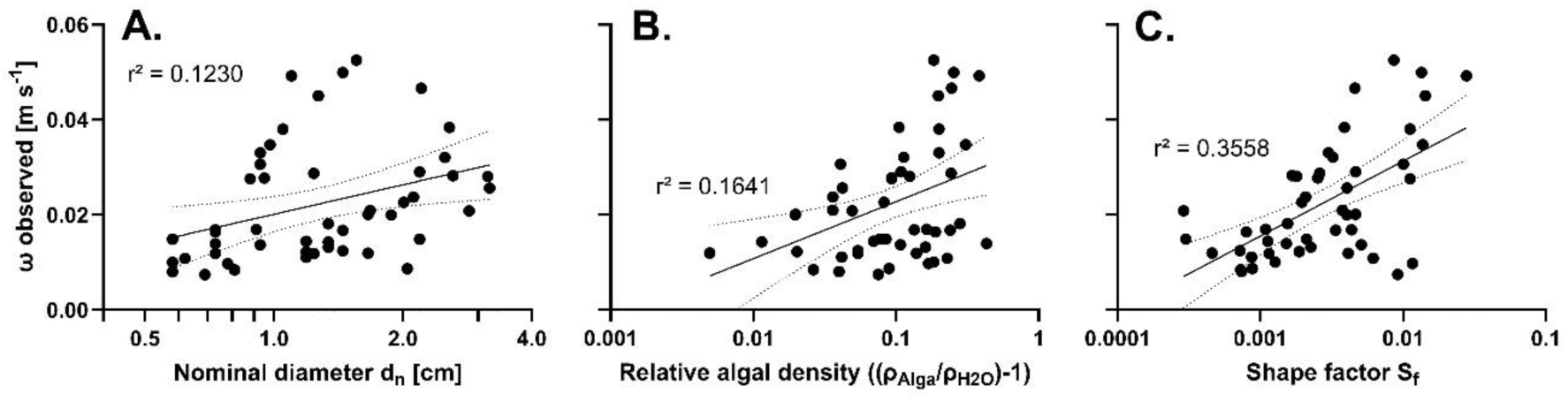
Correlations between sedimentation and morphological traits of macrophytes. Sedimentation velocity ω of 49 macrophyte specimens was correlated with (A) their nominal diameter, (B) their relative density and (C) their shape factor. Best fitting semilogarithmic functions and their 95 % confidence intervals are shown (in all three cases p < 0.0001).

### Modeling sinking velocity

Alltogether 20 macrophyte specimens (Tab. 1, “modeling” group) were used for the computation of models predicting sedimentation velocity. Resulting models were then tested for accuracy with the remaining 29 specimens (Tab. 1, “testing” group). The best-fitting sedimentation model that could be obtained without consideration of particle shape (S_f_) was

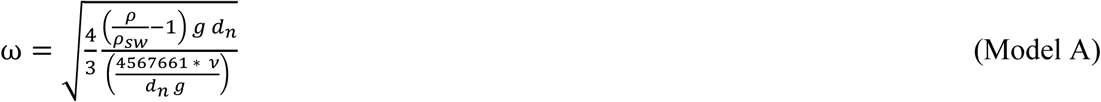

The sedimentation velocities predicted by this model correlated positively with the velocities observed, but the divergence between observed and predicted data was in many cases relatively large (Fig. 4A, r² = 0.3576, p < 0.0001). In particular, the mean sedimentation rate of the *S. latissima* rhizoid was significantly underestimated in the modeling dataset, placing it well outside the 95 % prediction interval. On the other hand, the velocity of specimen *D. sanguinea* 27 from the test sample set was overestimated. Alltogether, this model predicted the sinking velocities of the modeling sample set with a median squared deviation (MSD) of 6.84*10^−5^ and those of the test sample set with MSD = 4.28*10^−5^. Velocities of plastic disks were relatively accurately predicted, but velocities of nets underestimated and those of balls and rubber bands extremely underestimated (Fig. S2). As a consequence, the MSD of observed and predicted velocities of plastic items was relatively high (7.07*10^−4^).

**Fig. 4:**
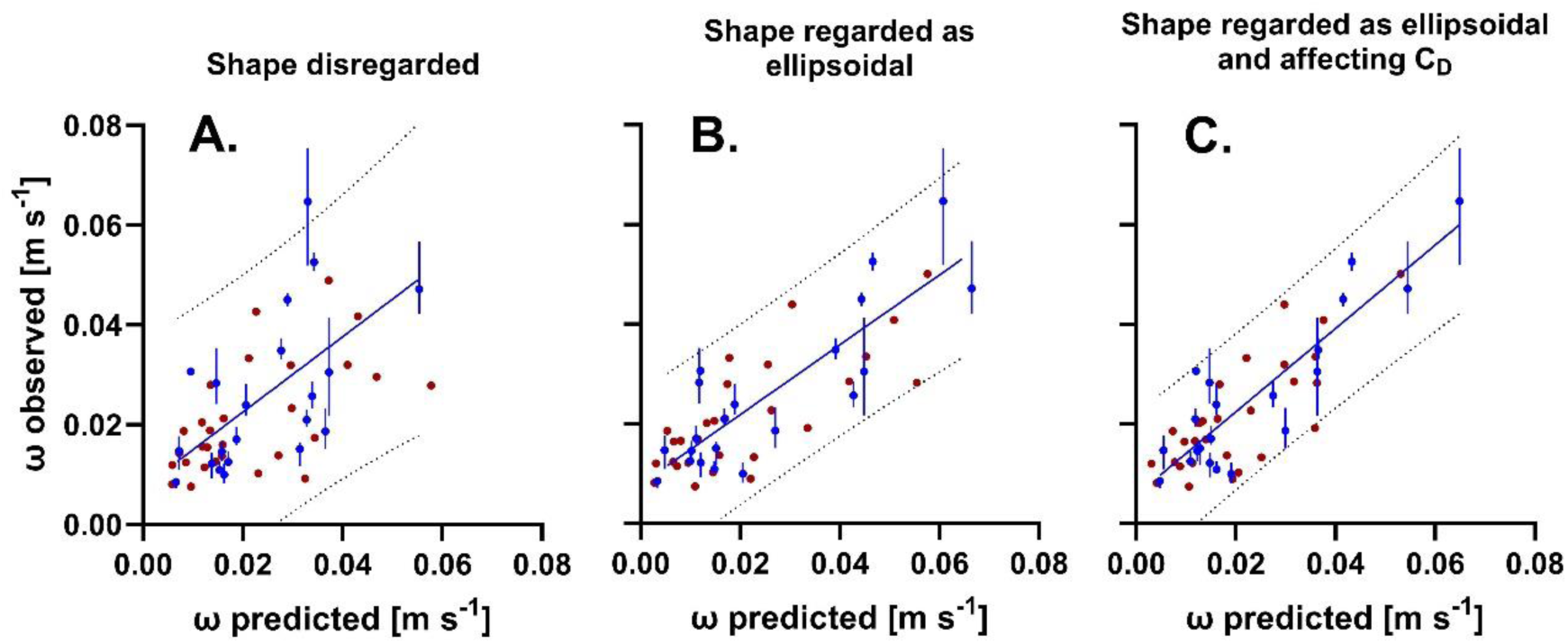
Correlations between observed and predicted sedimentation velocities. For the modeling sample set (blue, means ± ranges) and the test sample set (red, means only) with sedimentation velocities predicted for the same sample sets by (A) model A, (B) model B and (C) model C. Lines represent linear functions fitting best to data of the modeling sample set (model A: r² = 0.3576, p < 0.0001; model B: r² = 0.7210, p < 0.0001; model C: r² = 0.7926, p < 0.0001), dotted lines represent 95 % prediction intervals.

More prediction accuracy was possible when particle shape was considered in the model by inclusion of S_f_ into the numerator of the function and by taking into account the general ellipsoidal character of macrophytes. Under this condition the best fitting model that could be obtained was

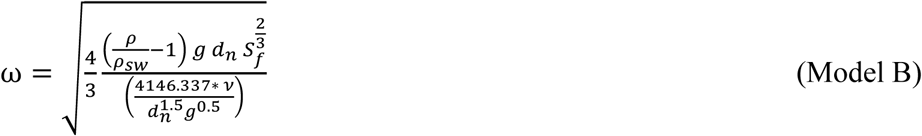

Model B (Fig. 4B, r² = 0.7210, p < 0.0001) visibly predicted the sedimentation velocities of macrophytes with more accuracy than model A (Fig. 4A). Correspondingly, MSDs obtained with model B (2.45*10^−5^ and 5.87*10^−5^ for the model sample set and the test sample set, respectively) were 29 % smaller than those obtained with model A. Model B also provided significantly better predictions of the sinking velocities of plastic items (Fig. S2B; MSD = 8.12*10^−5^).

The best prediction accuracy of the sinking velocity of macrophytes was achieved when S_f_ was not only included in the numerator, but also in the denominator of Eq. 4, considering that specific particle shape may also affect the drag coefficient C_D_. The best fitting model obtained under this condition was

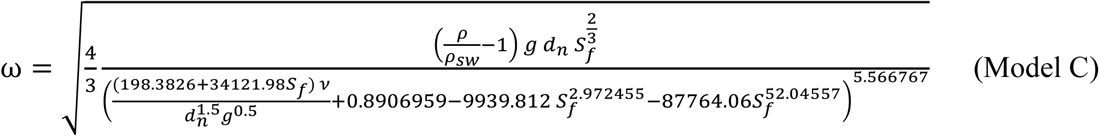

In particular the sedimentation velocities of faster sinking specimens were predicted with even more accuracy by model C (Fig. 4C, 0.7926, p < 0.0001) than by model B (Fig. 4B). Correspondingly, MSD’s obtained with model C (2.12*10^−5^ and 2.08*10^−5^ for modeling sample set and test sample set, respectively) were 40 % smaller than those obtained with model B. However, while model C predicted the sedimentation velocities of plastic discs approximately correctly it underestimated the velocities of nets (Fig. S2C). Moreover, model C extremely overestimated the sedimentation velocities of balls and rubber bands. As a consequence, the MSD of observed and predicted velocities of plastic items was only 8.00*10^−4^ and thus in the same order of magnitude as with model A, but 10 times larger than with model B.

Given that low S_f_ values characterised not only thin, leaf-shaped, but also filamentous, tufted algae (see Fig. 1), and since similar surface interactions with water are not *a priori* to be expected for such different morphologies, one might possibly expect less accurate predictions of ωby model C at low S_f_. However, macrophytes with lower S_f_ were not those with lower predicted sinking velocities and there was no significant correlation between S_f_ and and the MSD between predicted and observed velocities (Fig. S2A). Likewise, correlations between the relative algal density or the nominal diameter d_n_ and MSD were not observed (Figs. S2B and C).

In order to examine the interactive impact of the size and the specific shape of macrophytes on their sedimentation velocity more closely, fictitious values were entered for d_n_ and S_f_ in model C, resulting in the dependencies shown in Figure 5. Obviously, the influence of the shape factor increases with particle size. Large macrophytes with a small shape factor sink significantly more slowly than those with a large shape factor, while smaller macrophytes can generally be expected to have lower sinking velocities.

**Fig. 5:**
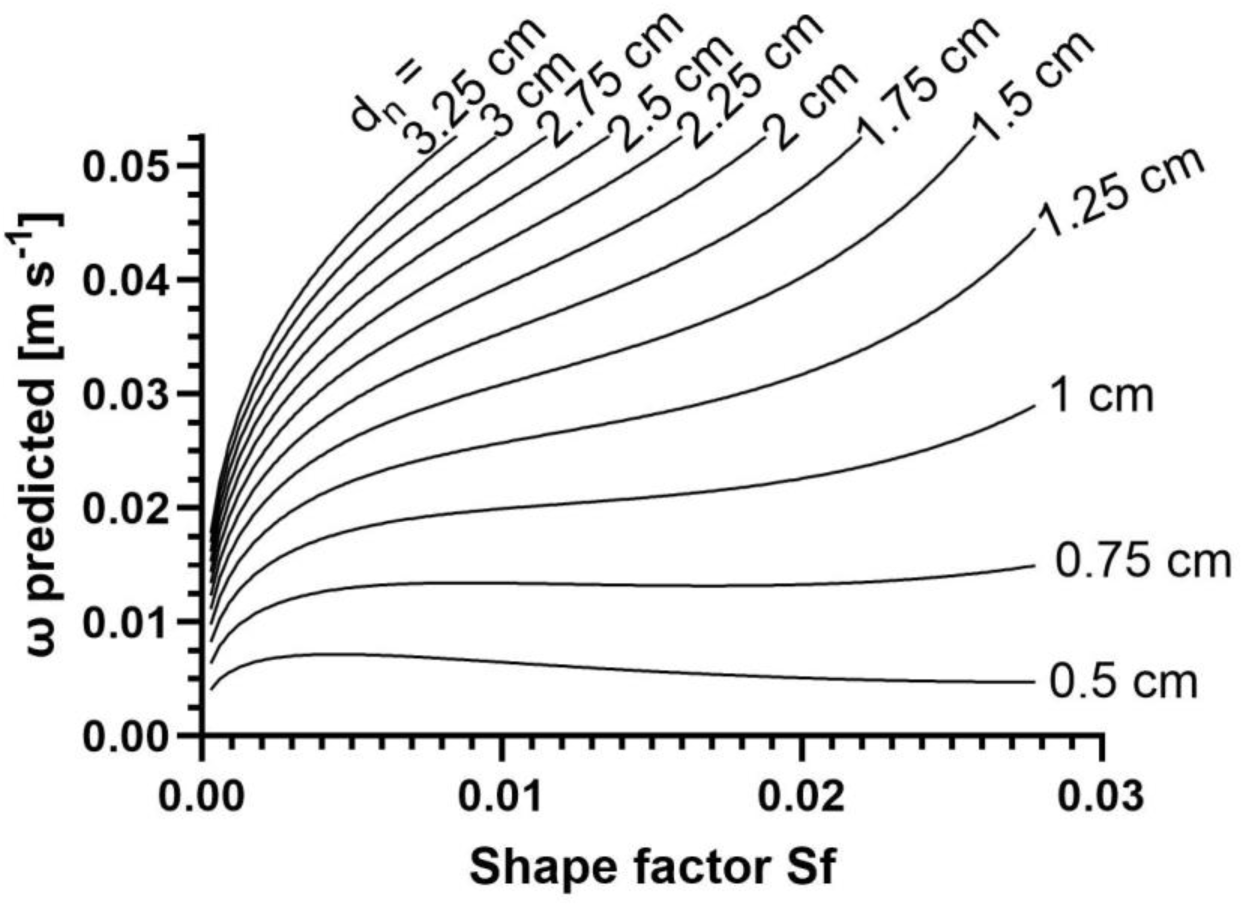
Predicted sinking velocity ω from model C. For macrophytes exhibiting different combinations of shape factor S_f_ and nominal diameter d_n_. Kinetic viscosity of sea water, macrophyte density and seawater density were considered to be the median conditions observed during our experimental setup, i.e. ρ = 1.0915*10^−6^ m^2^ s^−1^, ρ = 1104.44 kg m^−3^ and ρ_*H*2*O*_= 1009.45 kg m^−3^.

## Discussion

### Morphological differences in sedimentation behavior

This comparative study of morphologically very diverse marine macrophytes made it possible to identify both differences and similarities in their sedimentation behavior. With few exceptions (namely, *U. linza* and different thallus parts of *S. latissima*), specimens of the same species behaved similarly and we observed some tendencies toward different behaviour across major taxonomic groups. That is, brown and red seaweeds tended to sink faster than green seaweeds and flattened branched specimens faster than flattened unbranched ones.

### Predicting seaweed sinking velocity based on thallus volume, weight and shape

We identified the most relevant traits for the prediction of sinking velocity and drag impact on marine macrophytes and established protocols for their determination. The relevant parameters are (1) thallus volume, (2) thallus wet weight and (3) thallus shape. Thallus volume allows for the determination of the nominal diameter d_n_ as described in Eq. 5. Thallus volume and thallus wet weight together are required for the determination of thallus mass density ρ. Use of the Corey shape factor S_f_ proved to be an applicable solution to characterize thallus shape even in very irregularly shaped macrophytes. The assumption of an ellipsoidal particle shape is inherent to use of S_f_, given that S_f_ expresses the ratio of the longest, shortest and intermediate mutually perpendicular particle diameters. Leaving aside the more or less pronounced thallus flexibility, which can lead to considerable deviations, unbranched morphologies of macrophytes appear relatively obviously as ellipsoidal. A longest diameter is usually easy to identify in these cases. In the case of leaf-shaped morphologies the shortest diameter is the thallus thickness, in the case of cylindrical unbranched morphologies it is any mean diameter perpendicular to the longest axis. The ellipsoidal character of branched thallus morphologies, on the other hand, is less obvious. A longest diameter is often not clearly discernible and – where it becomes visible – in many cases still not easy to measure. In the case of side branches of varying length and thickness, there is also the question of how to determine a representative shortest diameter, not to mention a representative intermediate diameter. The method used in the present work, based on a direct measurement of the more obvious axis lengths (Table 1; a * b in the case of flattened morphologies, c in the case of cylindrical morphologies) and subsequent derivation of less obvious axis lengths (c in the case of flattened morphologies, a*b in the case of cylindrical morphologies) from the volume, proved to be practicable and led to convincing results.

The shape factors we found for macrophytes are in the range between 0.00029 and 0.069. A value of only 0.0003 was found with *Ulva gigantea*, which is characterised by particularly large, flat and very thin thalli. The individual we examined had a maximum length of 35 cm and a width of 29 cm with an average thallus thickness of only 79 µm. Individuals about twice or thrice this size exist (own observations) and a minimum value of S_f_ around 0.0001 could therefore be possible. Therefore, the minimum values observed in our study could be close to the actually existing minimum values of S_f_ in macrophytes. However, the maximum value of 0.069 is still well below the theoretically possible maximum value of 1, which would result for perfect spheres. It is to be expected that species such as *Codium bursa* or *Colpomenia peregrina*, which approximately have a spherical morphology but were not available for our study, will exceed the maximum value of S_f_ that we found.

### Enhancing accuracy by incorporating S_f_ as a component of C_D_

The inclusion of S_f_ values determined in this way into the drag equation enabled relatively accurate predictions of the sedimentation rate of macrophytes. Notably, the prediction accuracy increased when S_f_ was not only considered in the numerator, but also as a component of C_D_ in the denominator, as previously suggested by Riazi and Türker (2019) (Riazi and Türker 2019) for sediment particles. Remarkably, the best model C empirically determined on this premise and on the basis of our modeling data set predicted ω with slightly better accuracy for the test sample set than for the modeling data set.

### Differences in predicting the sedimentation velocity of plastic objects vs. seaweeds

In contrast, this model did not have good prediction accuracy for most plastic particles. Instead, it resulted in a significant overestimation of the sedimentation velocity for both balls and rubber bands. Only the velocities of the plastic disks were predicted relatively accurately, probably due to the fact that their shape factor is similar to that of macrophytes. Balls and rubber bands had shape factors that were significantly higher than those of all macrophytes. When balls and rubber bands were experimentally integrated into the modeling dataset, no alternative model based on the formula structure of model C could converge (not shown). Models based on this formula structure could therefore be fundamentally unsuitable for accurate predictions of the sedimentation velocity of particles with high shape factors. Alternatively, the very poor prediction accuracy of model C for balls and rubber bands could also result from the fact that the transition between laminar and turbulent conditions at the particle surface is influenced by the material properties (e.g., elasticity or compressibility) at the particle surface. These properties could differ between various plastic particles and macrophyte particles and were not considered further in our study. The observation that model C predicted the sinking speed of plastic nets with only moderate accuracy, although their form factors were in the same order of magnitude as those of macrophytes, also suggests an influence of surface material properties on C_D_.

### Model Performance divergence: C for seaweeds, B for plastic objects

On the other hand, the fact that model C worked with better accuracy than the simpler models A and B for all macrophytes tested indicates that the surface material properties of different macrophytes differ significantly less than their form factors in terms of influence on C_D_.

Interestingly, when the shape factor was only included in the numerator and not used to predict C_D_ (Model B), the sedimentation velocity of all plastic and macrophyte particles was predicted with reasonable accuracy and better accuracy than when shape was completely ignored (Model A). This demonstrates the general usefulness of considering S_f_.

## Conclusions

Our study highlights that the sedimentation behaviour and the sensitivity to drag of marine macrophytes and also of plastic particles can be predicted with significantly increased accuracy if they are regarded as ellipsoids and their specific shape is also considered. Model C performed well for a wide range of macrophyte species, which were characterised by very different morphologies. It can be assumed that the behaviour of macrophytes from other habitats – marine as well as limnic – would also be predicted with similar accuracy. With Model B, we were able to increase the predictive capacity for plastic particle shapes and sizes, although the model was less accurate than model C for macrophytes. Model C therefore appears as better suited for accurate predictions of the velocities of macrophytes, while Model B appears more robust for generalistic predictions, especially for particles with high shape factors. Potentially, our models could even provide sufficiently accurate estimates for other particles with sizes and specific densities similar to those of marine macrophytes, such as leaves shed by terrestrial plants. To accurately predict a higher variability of particles with higher shape factors than macrophytes, more diverse particles should be measured. A next step towards a generalistic model could also be to integrate different particle surface properties for a further model improvement.

Thus, our results provide the foundation for improving the accuracy of predictions of the sinking velocities of morphologically diverse seaweeds – an important step for understanding processes such as carbon sequestration and the delivery of biomass to deeper parts of the ocean. Additionally, they can help refine estimates of how fast negatively buoyant plastic objects transition from the surface to being deposited on the seafloor.

## Author contributions

FW, FG and TB initiated and designed this study. FG, FW and TB collected the data. FG and FW analysed the data. FW and FG generated the models. FW wrote the manuscript. All authors contributed to the final version of the manuscript.

## Data availability

Data available via the Pangaea Digital Repository (doi will be provided once paper is accepted).

## References

1. Barnes, D. K. A., Galgani, F., Thompson, R. C. and Barlaz, M. 2009. Accumulation and fragmentation of plastic debris in global environments. – Philos Trans R Soc Lond B Biol Sci 364: 1985–1998.

2. Bartlett, D. and Elmer, F. 2021. The impact of Sargassum inundations on the Turks and Caicos Islands. – Phycology 1: 83–104.

3. Bonsdorff, E. 1992. Drifting algae and zoobenthos — Effects on settling and community structure. – Netherlands Journal of Sea Research 30: 57–62.

4. Camenen, B. 2007. Simple and general formula for the settling velocity of particles. – J Hydraul Eng 133: 229–233.

5. Cheng, N. 1997. Simplified settling velocity formula for sediment particle. – J Hydraul Eng 123: 149–152.

6. Eriksen, M., Lebreton, L., Carson, H., Thiel, M., Moore, C., Borerro, J., Galgani, F., Ryan, P. and Reisser, J. 2014. Plastic Pollution in the World’s Oceans: More than 5 Trillion Plastic Pieces Weighing over 250,000 Tons Afloat at Sea. – PLoS ONE in press.

7. Galloway, T. S., Cole, M. and Lewis, C. 2017. Interactions of microplastic debris throughout the marine ecosystem. – Nat Ecol Evol 1: 1–8.

8. Garbossa, L. H. P., Santos, A. A. and Lapa, K. R. 2021. Seaweed dispersion under different environmental scenarios based on branches settling velocity and hydrodynamic lagrangian model. – Regional Studies in Marine Science in press.

9. Gewert, B., Plassmann, M. M. and MacLeod, M. 2015. Pathways for degradation of plastic polymers floating in the marine environment. – Environ. Sci.: Processes Impacts 17: 1513–1521.

10. Israel, A., Gavrieli, J., Glazer, A. and Friedlander, M. 2005. Utilization of flue gas from a power plant for tank cultivation of the red seaweed Gracilaria cornea. – Aquaculture (Amsterdam: 311–316.

11. Kokubu, Y., Rothäusler, E., Filippi, J.-B., Durieux, E. and Komatsu, T. 2019. Revealing the deposition of macrophytes transported offshore: Evidence of their long-distance dispersal and seasonal aggregation to the deep sea. – Scientific Reports in press.

12. Komar, P. D. and Reimers, C. E. 1978. Grain shape effects on settling rates. – J Geol 86: 193– 209.

13. Krause-Jensen, D. and Duarte, C. M. 2016. Substantial role of macroalgae in marine carbon sequestration. – Nature Geoscience 9: 737–742.

14. Kwon, K., Choi, B.-J., Kim, K. Y. and Kim, K. 2019. Tracing the trajectory of pelagic Sargassum using satellite monitoring and Lagrangian transport simulations in the East China Sea and Yellow Sea. – Algae 34: 315–326.

15. Mata, L., Santos, R., Chapman, A. R. O., Anderson, R. J., Vreeland, V. and Davison, I. R. 2003. Cultivation of Ulva rotundata (Ulvales, Chlorophyta) in raceways using semi-intensive fishpond effluents: yield and biofiltration.

16. Mossbauer, M., Haller, I., Dahlke, S. and Schernewski, G. 2012. Management of stranded eelgrass and macroalgae along the German Baltic coastline. – Ocean & Coastal Management 57: 1–9.

17. Msuya, F. E. and Neori, A. 2008. Effect of water aeration and nutrient load level on biomass yield, N uptake and protein content of the seaweed Ulva lactuca cultured in seawater tanks. – Journal of Applied Phycology 20: 1021–1031.

18. Nielsen, R., Lundsteen, S. and Brodie, J. 2023. Seaweeds of Denmark: vol. 1, Red algae (Rhodophyta) & vol. 2, Brown algae (Phaeophyceae) and Green algae (Chlorophyta). – Phycologia 62: 1–2.

19. Ortega, A., Geraldi, N. R., Alam, I., Kamau, A. A., Acinas, S. G., Logares, R., Gasol, J. M., Massana, R., Krause-Jensen, D. and Duarte, C. M. 2019. Important contribution of macroalgae to oceanic carbon sequestration. – Nature Geoscience 12: 748–754.

20. Oseen, C. W. 1927. Neuere Methoden und Ergebnisse in der Hydrodynamik. – Akademische Verlagsgesellschaft: 337.

21. Riazi, A. and Türker, U. 2019. The drag coefficient and settling velocity of natural sediment particles. – Computational Particle Mechanics 6: 427–437.

22. Rothäusler, E., Corell, H. and Jormalainen, V. 2015. Abundance and dispersal trajectories of floating Fucus vesiculosus in the Northern Baltic Sea. – Limnology and Oceanography 60: 2173–2184.

23. Sahoo, D. and Yarish, C. 2005. Mariculture of Seaweeds. – In: Andersen, R. A. (ed), Algal culturing techniques. Elsevier Academic Press, pp. 219–238.

24. Savvashe, P., Mhatre-Naik, A., Pillai, G., Palkar, J., Sathe, M., Pandit, R., Reddy, C. and Lali, A. 2021. High yield cultivation of marine macroalga Ulva lactuca in a multi-tubular airlift photobioreactor: A scalable model for quality feedstock. – Journal of Cleaner Production 329: 129746.

25. Schindelin, J. 2012. Fiji: an open-source platform for biological-image analysis. – Nature methods 9: 676–682.

26. Scrucca, L. 2013. GA: A Package for Genetic Algorithms in R. – Journal of Statistical Software 53: 1–37.

27. Sharqawy, M. H., Lienhard, J. H. and Zubair, S. M. 2012. Thermophysical properties of seawater: a review of existing correlations and data. – Desalination and Water Treatment 16: 354–380.

28. She, K., Trim, L. and Pope, D. H. 2005. Fall velocities of natural sediment particles: a simple mathematical presentation of the fall velocity law. – J Hydraul Res 43: 189–195.

29. Smetacek, V. and Zingone, A. 2013. Green and golden seaweed tides on the rise. – Nature 504: 84–88.

30. Stokes, G. G. 1851. On the effect of the internal friction of fluids on the motion of pendulums. – Transactions of the Cambridge Philosophical Society: 8.

31. Swamee, P. and Ojha, C. 1991. Drag coefficient and fall velocity of nonspherical particles. – J Hydraul Eng 117: 660–667.

32. Vahteri, P., Mäkinen, A., Salovius, S. and Vuorinen, I. 2000. Are Drifting Algal Mats Conquering the Bottom of the Archipelago Sea, SW Finland? -Ambio (Sweden) 29: 338–343.

33. Wadell, H. 1935. Volume, shape, and roundness of quartz particles. – Journal of Geology in press.

34. Weinberger, F., Buchholz, B., Karez, R. and Wahl, M. 2008. The invasive red alga Gracilaria vermiculophylla in the Baltic Sea: adaptation to brackish water may compensate for light limitation. – Aquatic Biology 3: 251–264.

35. Weinberger, F., Paalme, T. and Wikström, S. A. 2020. Seaweed resources of the Baltic Sea, Kattegat and German and Danish North Sea coasts. – Botanica Marina 63: 61–72.

36. Weinberger, F., Sundt, S., Staerck, N., Merk, C., Karez, R. and Rehdanz, K. 2021. Shifting beach wrack composition in the SW Baltic Sea and its effect on beach use. – Ecology and Society in press.

37. Zhang, J. 2019. Annual patterns of macroalgal blooms in the Yellow Sea during 2007-2017. – PloS one 14: 0210460.

38. Zhou, F., Ge, J., Liu, D., Ding, P., Chen, C. and Wei, X. 2021. The Lagrangian-based Floating Macroalgal Growth and Drift Model (FMGDM v1.0): application to the Yellow Sea green tide. – Geoscientific Model Development 14: 6049–6070.

