## Supplementary material for "Sedimentation and drag in drifting macrophytes and plastic objects: A model"

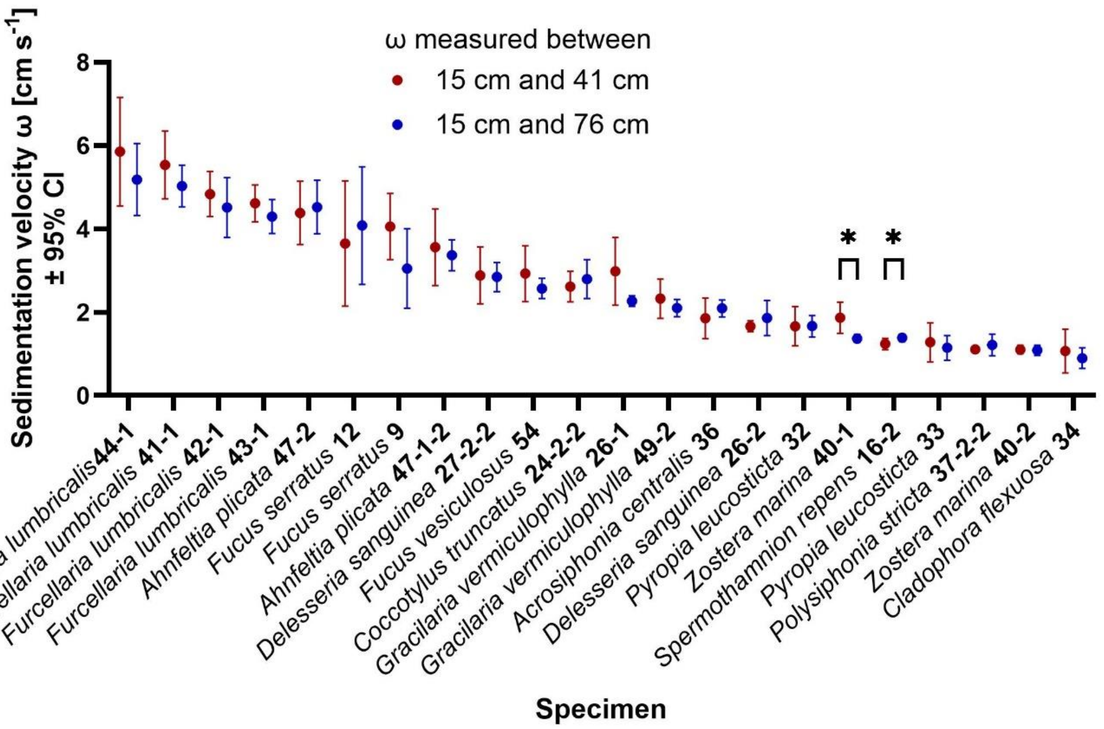


**Fig S1: Comparison of sinking velocities of different macrophytes measured with two different methods.** The specimens were released at the water surface and their velocity was measured after they had reached a depth of 15 cm. One approach measured ω in the water depth range between 15 cm and 41 cm, while the second measured it in the range between 15 cm and 76 cm. Asterisks indicate specimens for which significantly different results were obtained with the two approaches (Welch-corrected t-test, n = 5, $p < 0.05$).

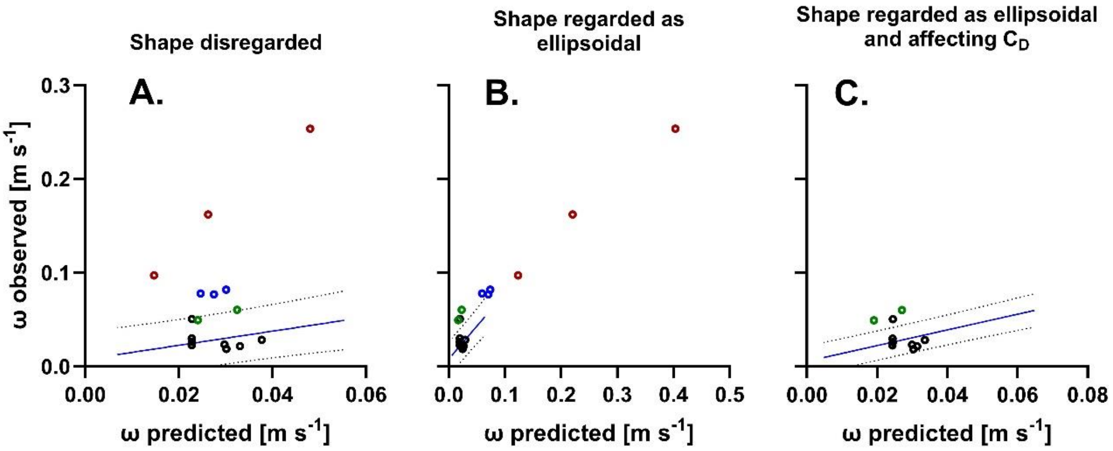


**Fig. S2: Correlations of sedimentation velocities observed for plastic particles.** Discs (black), nets (green), rubber bands (blue) and balls (red) with sedimentation velocities predicted for the same sample sets by (A) model A, (B) model B and (C) model C. In C data for balls and rubber bands are not shown, as predictions were extremely high (> $10^{99}$ m $s^{-1}$). Lines represent the linear functions that fitted best to data of the modeling macrophyte sample set (see also Figure 4), dotted lines represent 95 % prediction intervals.

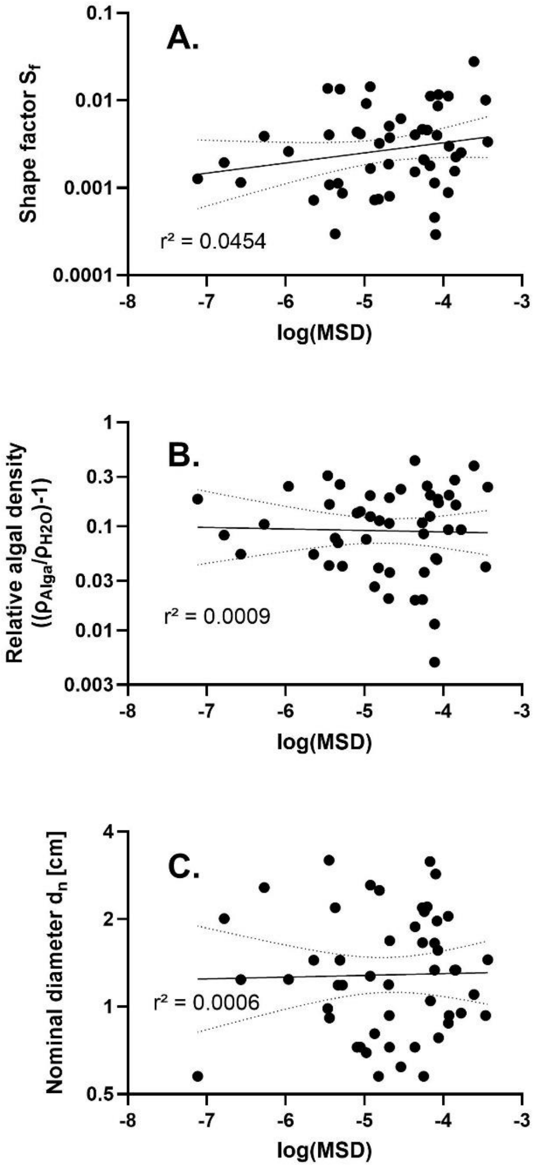


**Fig S3: Double logarithmic correlations of morphological traits.** (A) the shape factor, (B) the relative mass density and (C) the nominal diameter of 49 macrophyte specimens with the mean square deviation (MSD) of sinking velocities predicted by model C and observed with these specimens. Best fitting functions and their 95 % confidence intervals are shown (in all three cases $p > 0.05$).

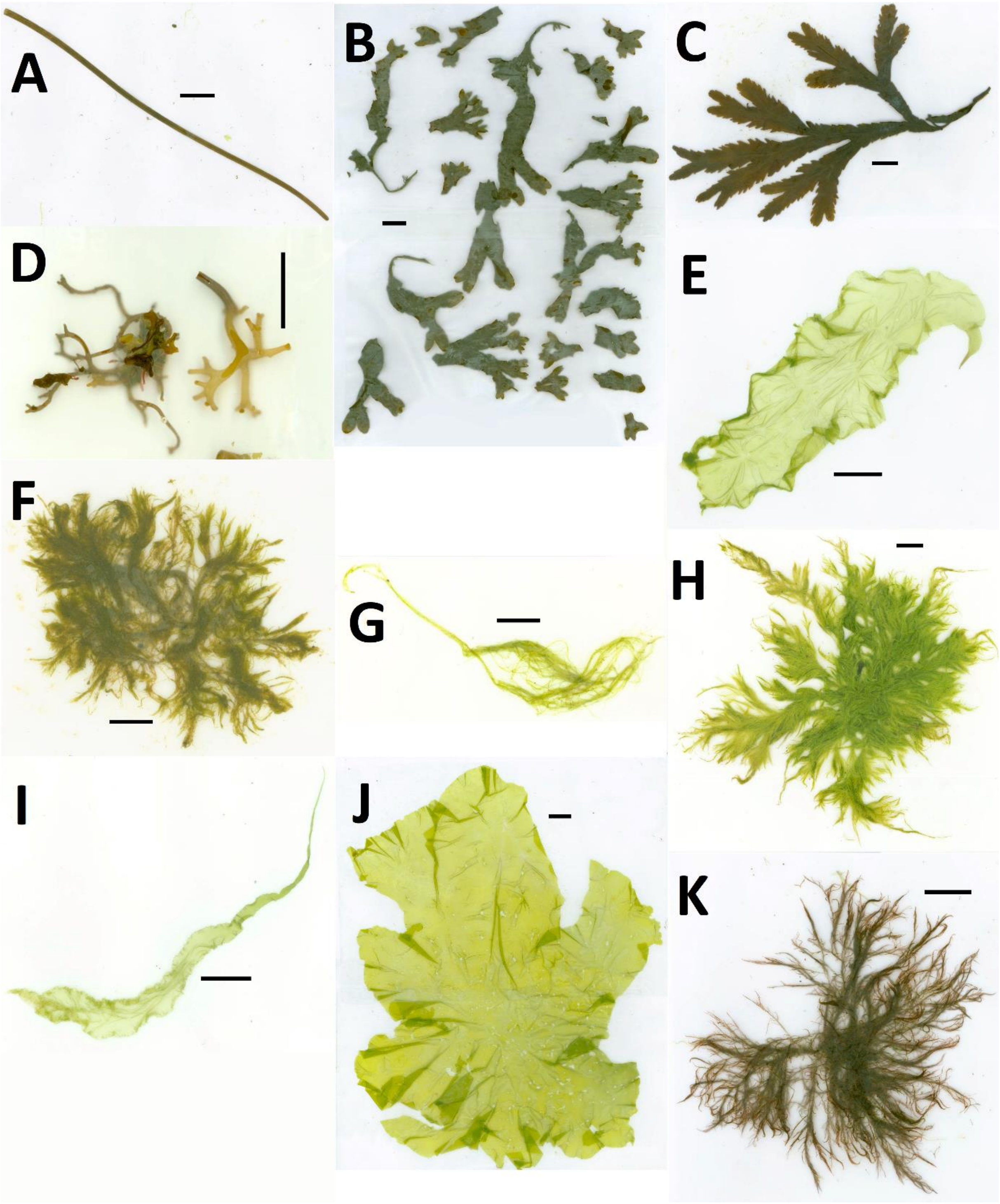


**Fig. S4: Specimens of the modeling sample set.** (A) *Z. marina* 40-2, (B) *F. vesiculosus* 54, (C) *F. serratus* 9, (D) *S. latissima* rhizoid K9B, (E) *K. leptoderma* 1-1, (F) *B. hypnoides* 1-2, (G) *U. clathrata* 21-1, (H) *A. centralis* 36, (I) *U. linza* 8-1, (J) *U. gigantea* 10, (K) *P. stricta* 37-2-1. Horizontal or vertical scale bars represent a distance of 2 cm, Specimens in B and D were disassembled to facilitate scanning.

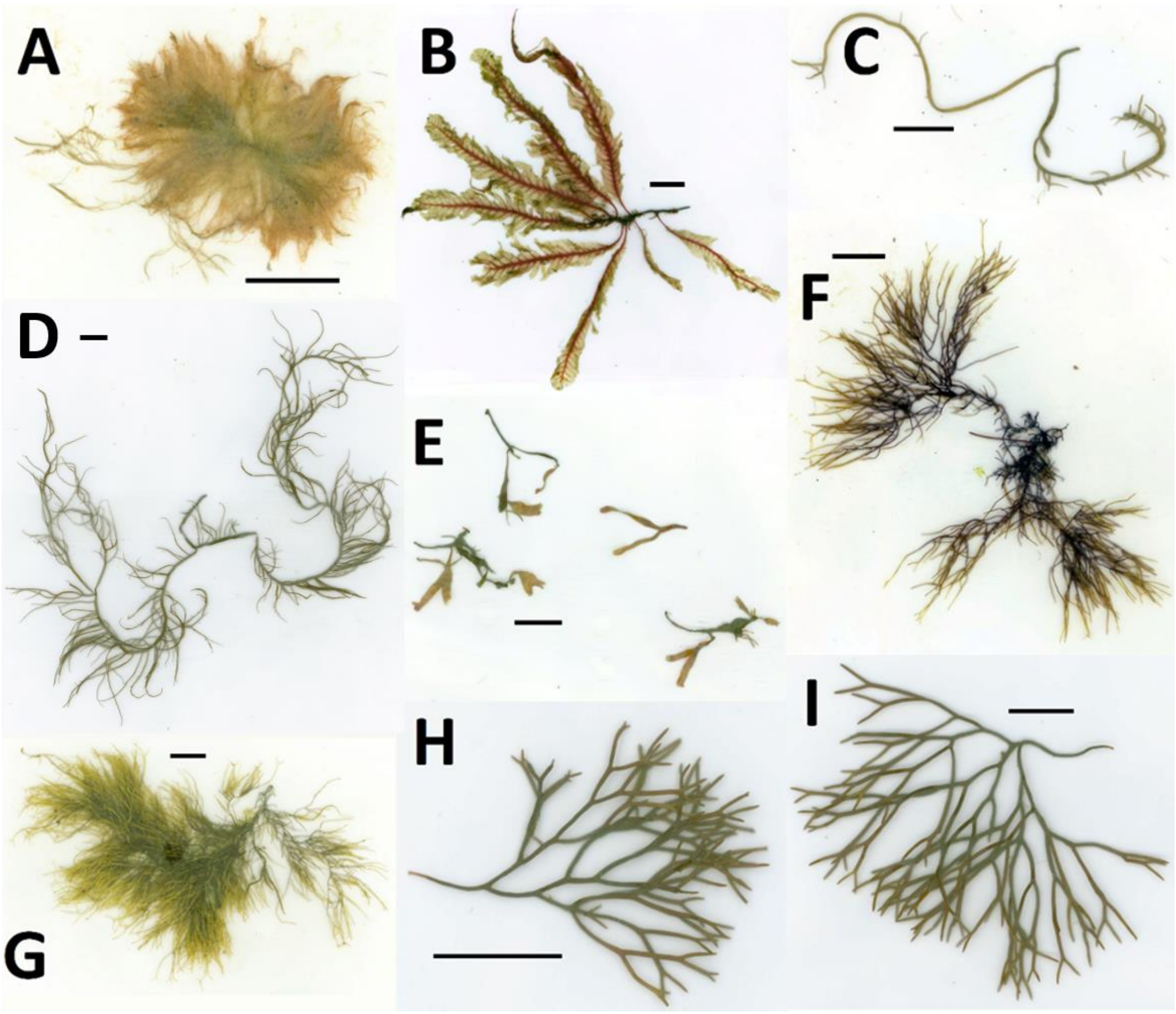


**Fig. S5: Specimens of the modeling sample set.** (A) *S. repens* 14, (B) *D. sanguinea* 26-2, (C) *G. vermiculophylla* 16-1, (D) *G. vermiculophylla* 49 (E) *C. truncatus* 24-2-1, (F) *A. plicata* 47-1, (G) *A. plicata* 47-2, (H) *F. lumbricalis* 43-1, (I) *F. lumbricalis* 44-1. Horizontal scale bars represent a distance of 2 cm. The specimen in E was disassembled to facilitate scanning.

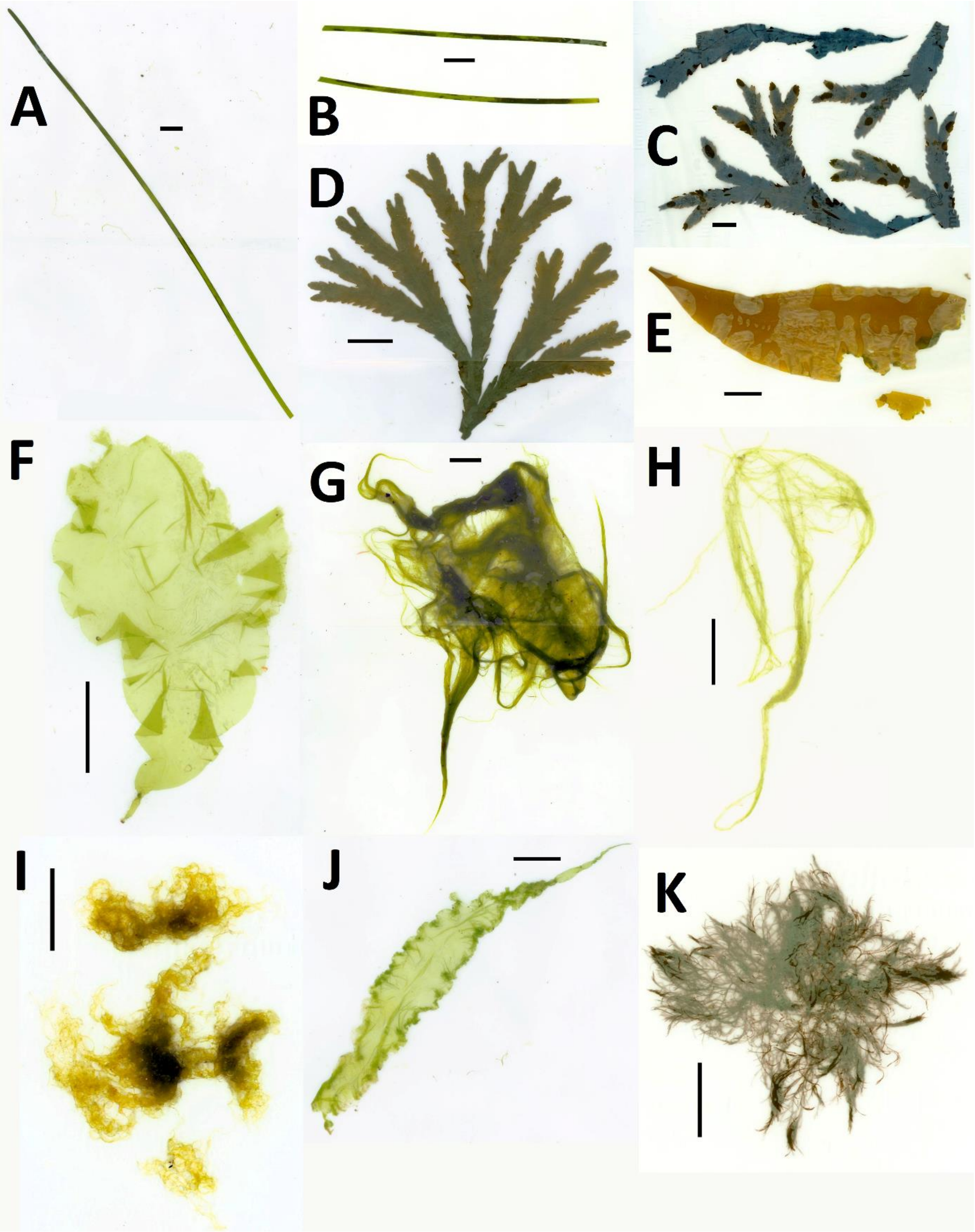


**Fig. S6: Specimens of the testing sample set.** (A) *Z. marina* 40-1, (B) *Z. marina* K7, (C) *F. serratus* K5, (D) *F. serratus* 12, (E) *S. latissima* phylloid K9A, (F) *K. leptoderma* 4-2, (G) *C. flexuosa* 34, (H) *U. clathrata* 24-1, (I) *Cladophora* sp. K3, (J) *U. linza* 6-1, (K) *V. fucoides* K2. Horizontal or vertical scale bars represent a distance of 2 cm. Specimens in B, C and I were disassembled to facilitate scanning, specimen E broke prior to scanning.

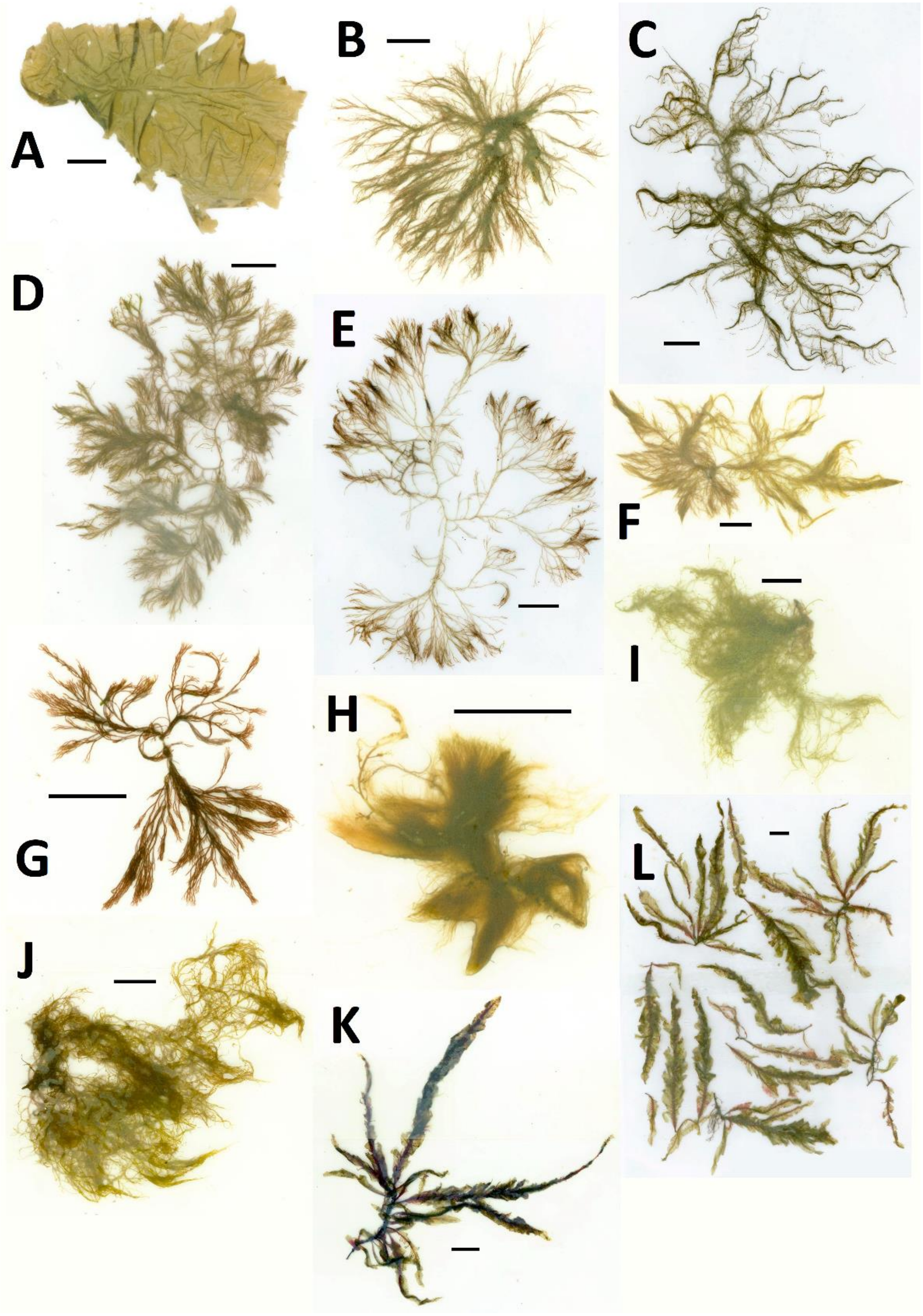


**Fig. S7: Specimens of the testing sample set.** (A) *P. leucosticta* 32, (B) *P. stricta* 37-2-2, (C) *P. stricta* 39-2, (D) *C. virgatum* 6-2, (E) *C. virgatum* 8-2, (F) *C. virgatum* 7-2, (G) *C. virgatum* K6, (H) *S. repens* 16-2, (I) *R. confervoides* 4-1-2, (J) *R. confervoides* 4-1, (K) *D. sanguinea* 27, (L) *D. sanguinea* 27-2-2. Horizontal or vertical scale bars represent a distance of 2 cm. The specimen in L was disassembled to facilitate scanning.

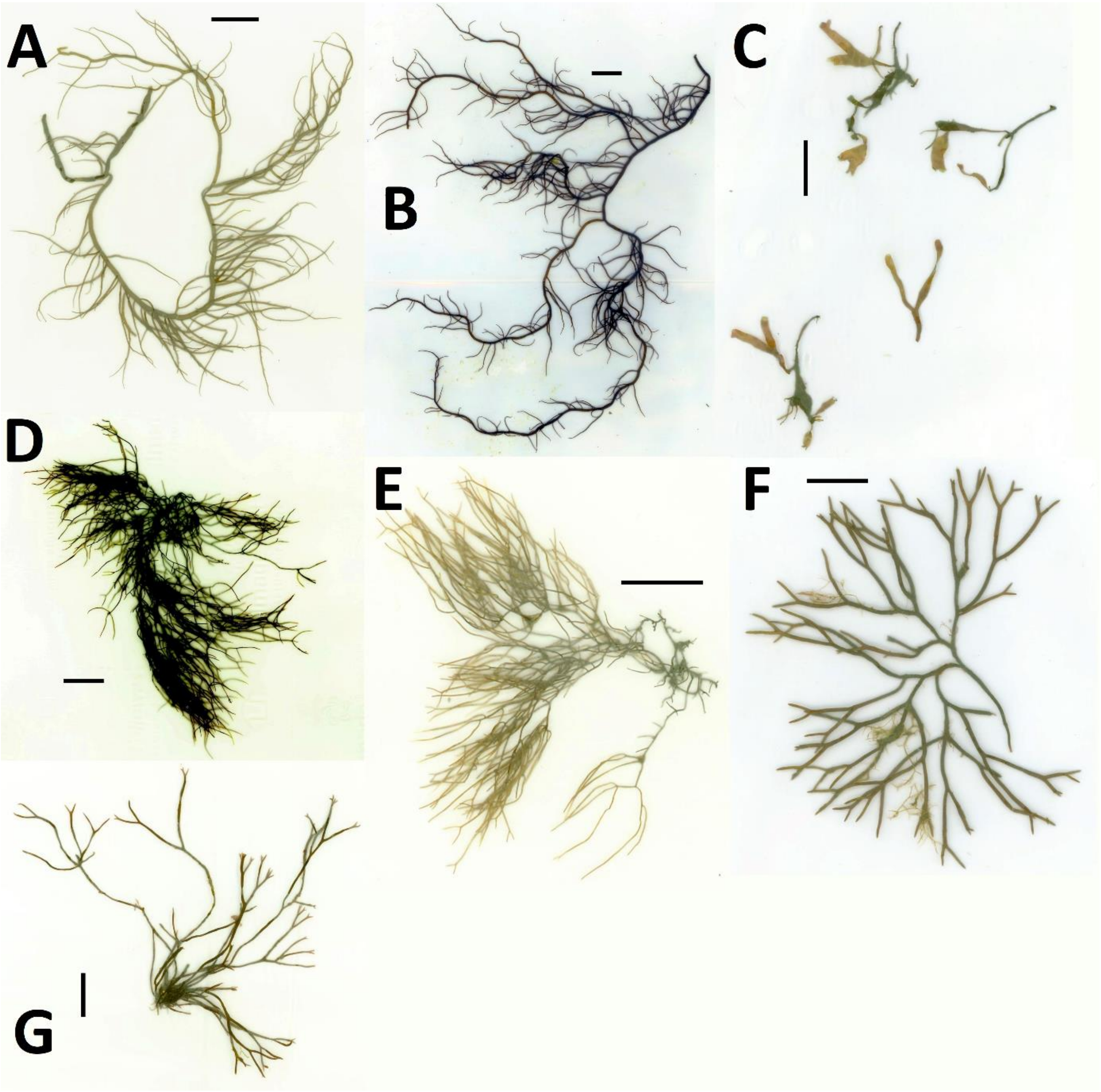


**Fig. S8: Specimens of the testing sample set.** (A) *G. vermiculophylla* 49-2, (B) *G. vermiculophylla* 26-1, (C) *C. truncatus* 24-2-2, (D) *A. plicata* K1, (E) *A. plicata* 47-1-2, (F) *F. lumbricalis* 41-1, (G) *F. lumbricalis* K8. Horizontal or vertical scale bars represent a distance of 2 cm. The specimen in C was disassembled to facilitate scanning.

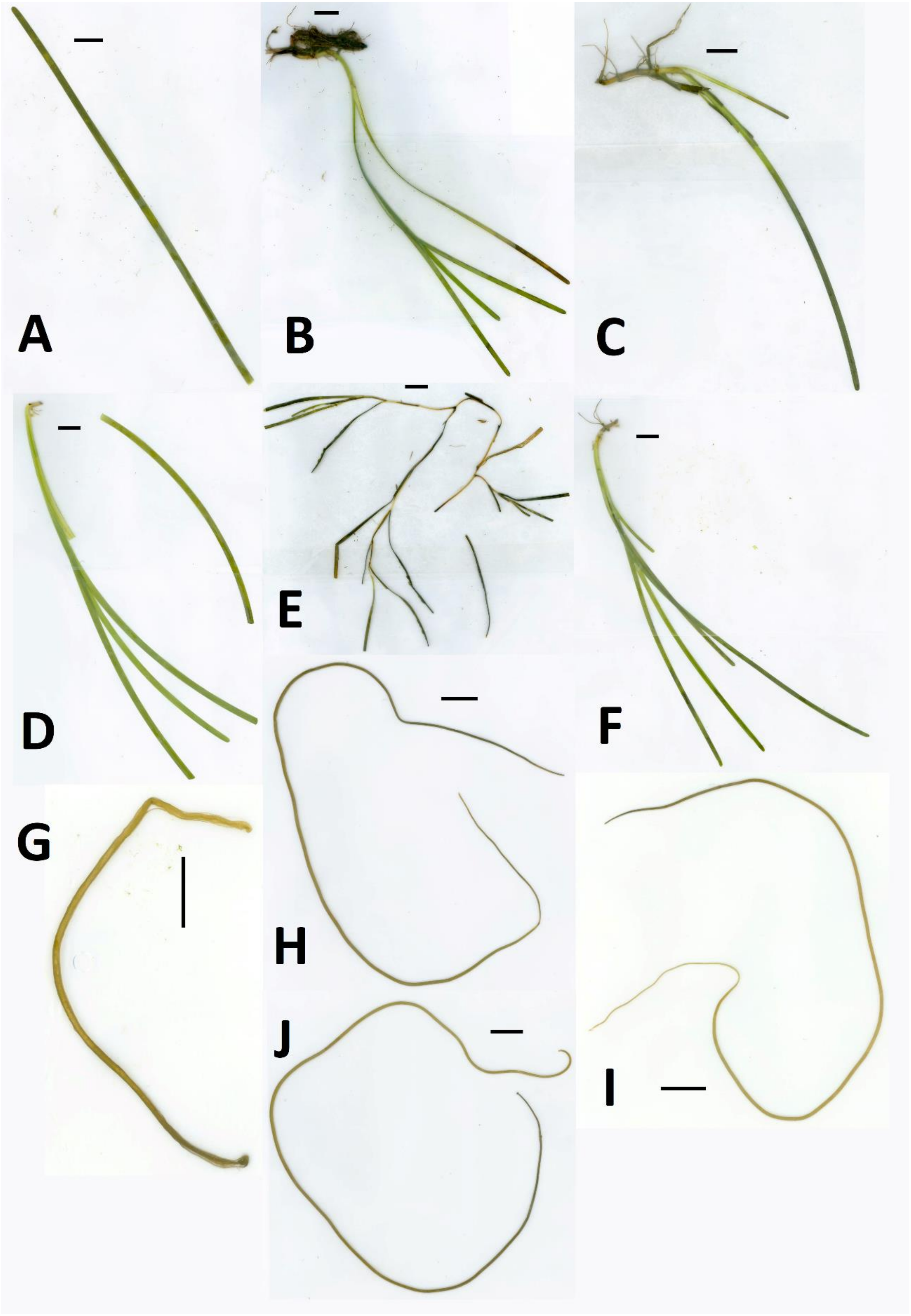


**Fig. S9: Positively buoyant specimens.** (A) *Z. marina* 39-1, (B) *Z. marina* 37-1, (C) *Z. marina* 69, (D) *Z. marina* K1, (E) *Z. marina* 45, (F) *Z. marina* 48-1, (G) *C. filum* 41-2, (H) *C. filum* 44-2, (I) *C. filum* 43-2, (J) *C. filum* 24-2. Horizontal or vertical scale bars represent a distance of 2 cm. Specimens in D and E broke prior to scanning.

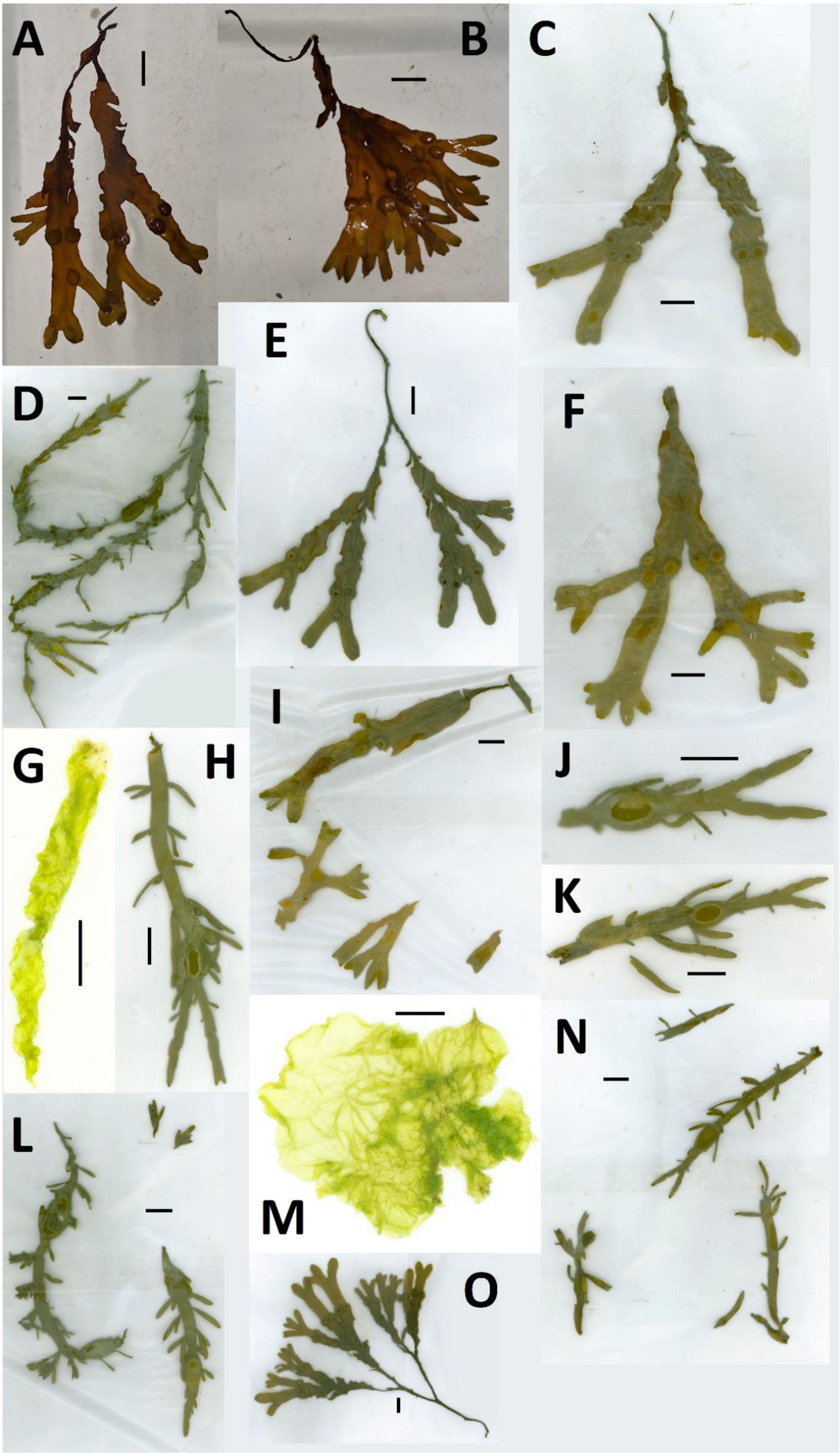
A
B
C
D
E
F
G
H
I
J
K
L
M
N
O

**Fig. S10 (previous page): Positively buoyant specimens.** (A) *F. vesiculosus* 3, (B) *F. vesiculosus* 35, (C) *F. vesiculosus* 38, (D) *A. nodosum* 29, (E) *F. vesiculosus* 15, (F) *F. vesiculosus* 13, (G) *U. linza* 7-1, (H) *A. nodosum* 5, (I) *F. vesiculosus* 51, (J) *A. nodosum* 46, (K) *A. nodosum* 2, (L) *A. nodosum* 68, (M) *U. compressa* 48-2, (N) *A. nodosum* 21, (O) *F. vesiculosus* 53. Horizontal or vertical scale bars represent a distance of 2 cm. Specimens in I, L and N were disassembled to facilitate scanning.

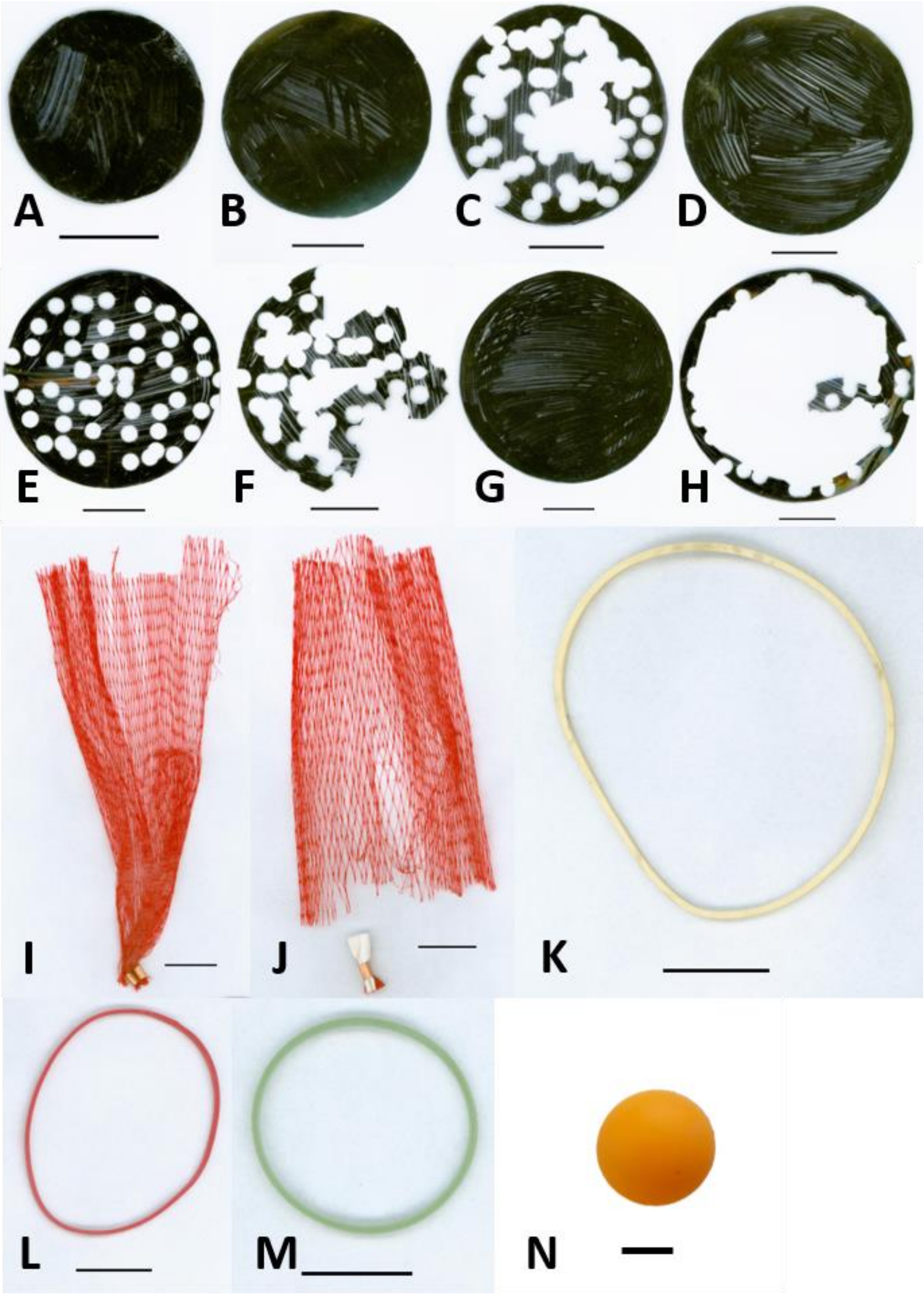


**Fig. S11: Plastic particles.** (A) Disc 40-1, (B) Disc 59-1, (C) Disc 59-2, (D) Disc 70-1, (E) Disc 70-2, (F) Disc 70-3, (G) Disc 84-1, (H) Disc 84-4, (I) Net-large, (J) Net-small, (K) Rubberband-large, (L) Rubberband-medium, (M) Rubberband-small, (N) Ball 1, 2, 3. Horizontal or vertical scale bars represent a distance of 2 cm.
